# Antibodies disrupt bacterial adhesion by ligand mimicry and allosteric interference

**DOI:** 10.1101/2024.12.06.627246

**Authors:** Kelli L. Hvorecny, Gianluca Interlandi, Tim S. Veth, Pavel Aprikian, Anna Manchenko, Veronika L. Tchesnokova, Miles S. Dickinson, Joel D. Quispe, Nicholas M. Riley, Rachel E. Klevit, Pearl Magala, Evgeni V. Sokurenko, Justin M. Kollman

**Affiliations:** Department of Biochemistry, University of Washington, Seattle, WA; Department of Bioengineering, University of Washington, Seattle, WA; Department of Chemistry, University of Washington, Seattle, WA; Department of Microbiology, University of Washington, Seattle, WA

## Abstract

A critical step in infections is the attachment of many microorganisms to host cells using lectins that bind surface glycans, making lectins promising antimicrobial targets. Upon binding mannosylated glycans, FimH, the most studied lectin adhesin of type 1 fimbriae in *E. coli*, undergoes an allosteric transition from an inactive to an active conformation that can act as a catch-bond. Monoclonal antibodies that alter FimH glycan binding in various ways are available, but the mechanisms of these antibodies remain unclear. Here, we use cryoEM, mass spectrometry, binding assays, and molecular dynamics simulations to determine the structure-function relationships underlying antibody-FimH binding. Our study reveals four distinct antibody mechanisms of action: ligand mimicry by an N-linked, high-mannose glycan; stabilization of the ligand pocket in the inactive state; conformational trapping of the active and inactive states; and locking of the ligand pocket through long-range allosteric effects. These structures reveal multiple mechanisms of antibody responses to an allosteric protein and provide blueprints for new antimicrobial that target adhesins.

## Introduction

Microbes adhere to surfaces as the first step in the infection of a host and in the formation of biofilms on both biological and abiotic surfaces ^1^. Adhesion also facilitates the mutualistic relationships found in the gastrointestinal tracts of various species ^2^. Many viruses, bacteria, fungi, and parasites use lectin-based, cell attachment proteins during infection or surface adherence, recognizing specific moieties on linear or branched oligosaccharide chains on the target surfaces ^3^.

FimH, from *Escherichia coli*, is the best characterized member of the largest family of structurally diverse fimbrial adhesins found in gram-negative bacteria, which are assembled via the chaperone-usher pathway^4^ (**Ext. Data Fig1a-b**). In *E. coli*, FimH acts as a key virulence factor in urinary tract infections and inflammatory bowel disease. FimH is located at the tip of the rod-shaped, type 1 fimbriae (pili) and contains two immunoglobulin-like domains: the lectin domain, which binds α-D-linked mannoses of N-linked glycans; and the pilin domain, which connects the lectin domain to additional subunits on the fimbria ^5–9^.

FimH also serves as a model allosteric protein. Upon ligand binding, FimH undergoes dramatic conformational rearrangements ^7,10^. In the absence of ligand, FimH adopts a compressed and inactive conformation stabilized by a fixed interface between the lectin and pilin domains, with the ligand-binding pocket relaxed and open ^10^ (**Ext. Data Fig. 1c**). Ligand binding closes the pocket and shifts FimH into an intermediate conformation. In the intermediate conformation, the mannose-binding loops, N-terminus, and clamp loop encircle the ligand, but the lectin-pilin domain interface remains fixed ^11^ (**Ext. Data Fig. 1d**). The core conformational changes caused by ligand binding propagate through the lectin domain and increase the likelihood that the lectin and pilin domains will separate and adopt the elongated active conformation ^11,12^ (**Ext. Data Fig. 1e**). Tensile mechanical force facilitates lectin-pilin domain separation, which pulls the pilin domain away from the lectin domain, stabilizing the active conformation and resulting in a force-enhanced catch-bond interaction ^7,10,13,14^.

**Figure 1:**
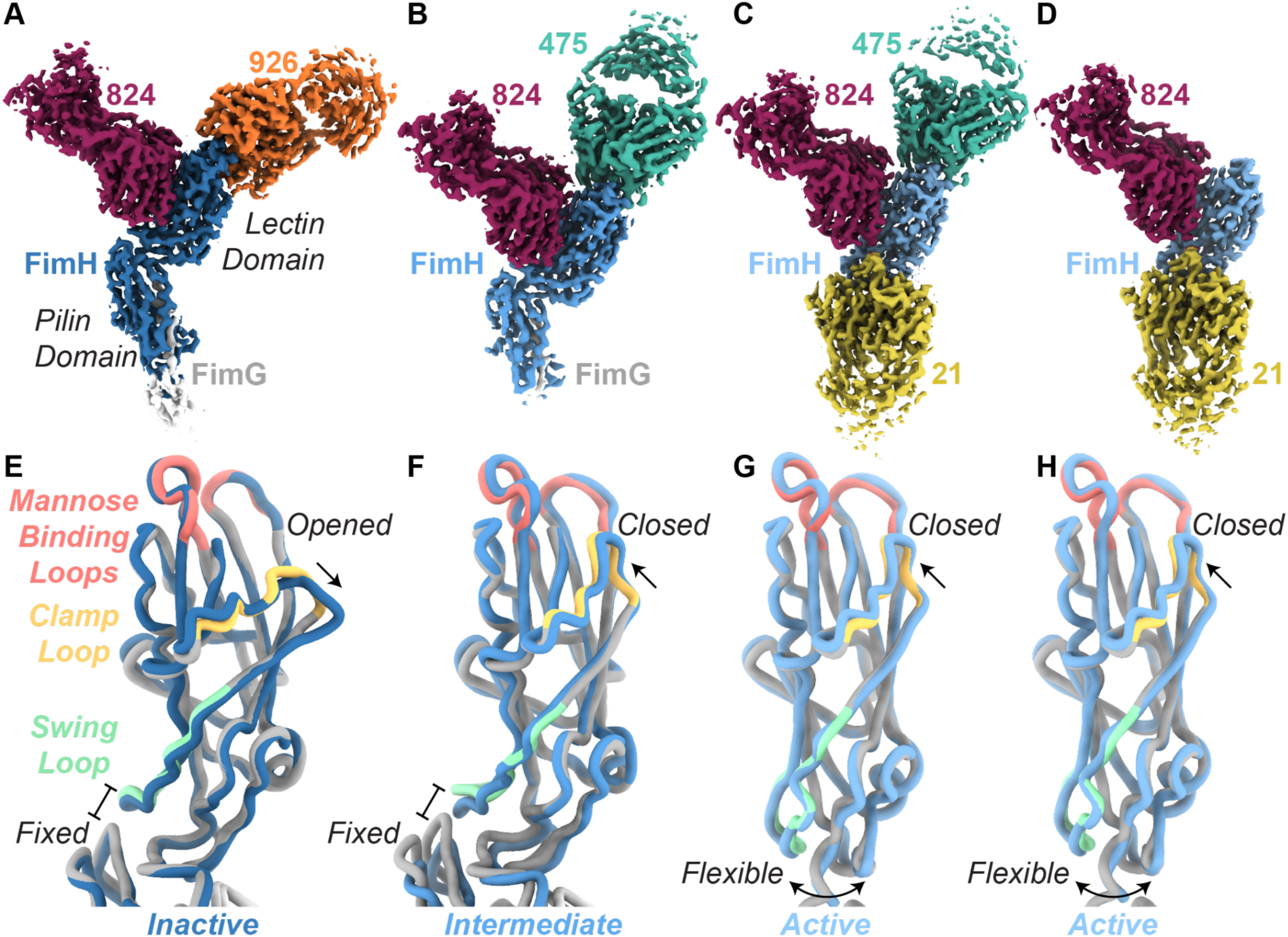
FimH conformations from fimbrial tip-Fab structures. **A-D**. Reconstructions of FimH bound to: **A**, Fab824 (maroon) and Fab926 (orange); **B**, Fab475 (teal) and Fab824 (maroon); **C**, Fab21 (gold), Fab475 (teal), and Fab824 (maroon); **D**. Fab21 (gold), Fab824 (maroon), and α-methyl-mannose (out of view). **E-H**. Alignment of FimH models from this work (blue) with previously published crystal structures (grey; swing loop in pale green, clamp loop in pale yellow, and mannose binding loops in pale red). **E**. Ribbon diagram of FimH-Fab824/926 aligned to PDB ID 3JWN. **F**. Ribbon diagram of FimH-Fab475/824 aligned to PDB ID 4XOE. **G-H**. Ribbon diagrams of FimH-Fab21/475/824 and FimH-Fab21/824/α-methyl-mannose aligned to PDB ID 1KLF.

Because of the critical role adhesins play as virulence factors in infection and biofilm formation, adhesins are attractive antimicrobial targets. Several strategies for inhibition of adhesins have been explored. For example, researchers have created compounds containing mannose and other glycan motifs to competitively inhibit surface glycan binding, a strategy similar to mammalian secretion of mucins and oligosaccharides to prevent pathogen binding ^15–22^. Researchers have also used FimH and other adhesins as vaccine antigens to elicit neutralizing antibodies ^23–25^. Follow-up studies, however, demonstrate that the polyclonal antibody response is complex and varies depending on the specific form of FimH used for vaccination ^11,23,26,27^.

Our earlier studies identified monoclonal antibodies (mAb) that impact the allosteric properties and neutralize FimH via distinct and previously not appreciated mechanisms ^28^. We classified the monoclonal antibodies into four categories: orthosteric, parasteric, dynasteric, and activating. *Ortho*steric antibodies, like mAb475, directly compete for the ligand binding pocket and behave like classical competitive inhibitors ^26^. mAb475 acts similarly to FimH ligands; the core epitope includes all mannose binding residues on FimH and the antibody triggers the transition from the inactive conformation to the active conformation (**Ext. Data Fig. 1f**). In contrast, *para*steric antibodies (mAb926) exhibit non-competitive behavior and the core epitope includes a few mannose-binding residues as well as nearby residues ^26^ (**Ext. Data Fig. 1f**). mAb926 stabilizes the inactive conformation of FimH and the antibody can uniquely elute bacteria already attached to a mannose-coated surface. *Dyna*steric antibodies, (mAb824), arrest the conformational dynamics of FimH; the antibodies bind both active and inactive conformations of FimH, but prevent the transition between conformations ^27^. The core epitope for mAb824 is distal to the ligand-binding pocket (**Ext. Data Fig. 1f-g**). Activating antibodies, like mAb21, only bind the active conformation of FimH ^23,26,27^. The core epitope for mAb21 includes residues in the interdomain region of the lectin domain (**Ext. Data Fig. 1f**). While our previous studies provided insights into the binding epitopes and large-scale conformational rearrangements that occur upon antibody binding, we lack the understanding of how each antibody exerts its effect at the molecular level, information which is required for the development of novel therapies against FimH and other cell attachment proteins of human pathogens.

Here, we investigate the antibody mechanisms of action of four functionally distinct antibodies. The orthosteric mAb475 leverages an unexpected N-linked, high-mannose glycan on a hypervariable loop to mimic the native, glycosylated host cell receptors to competitively block adhesion. The parasteric mAb926 occupies the open ligand-binding pocket yet retains partial flexibility in the antibody hypervariable loops which generates non-competitive inhibition. The dynasteric mAb824 directly engages allosteric residues on FimH to block conformational transitions. The activating mAb21 acts as a molecular wedge between the lectin and pilin domains, while allowing FimH to retain flexibility at the lectin-pilin interface. Each FimH-antibody interaction provides mechanistic insights with implications for the development of anti-adhesin therapies, with broad implications for human responses to the cell-binding proteins of pathogens.

## Results

### CryoEM structures reveal binding modes and FimH conformations

We used single-particle cryoEM to determine structures of fimbrial tips in complex with antibody fragments (Fabs) (**Ext. Data Fig. 2-4**, **Table 1**). Full fimbrial tips (FimH/G/F with chaperone FimC) complexed with multiple Fabs without overlapping epitopes increased the size of our target while allowing us to unambiguously identify each Fab bound to FimH (**Figure 1a-d, Ext. Data Fig. 5a**). We could not resolve components of the tip beyond FimG, likely due to flexibility (**Ext. Data Fig. 3**). Both Fab926 and Fab475 bind the mannose pocket of FimH but in distinct orientations relative to FimH (**Figure 1a-c, Ext. Data Fig. 5a**). Fab824 binds to the side of the lectin domain (**Figure 1a-d).** Fab21 interacts with the lectin domain at the lectin-pilin domain interface (**Figure 1c-d, Ext. Data Fig. 5a**). This agrees with previous mutational analyses that identified the functional epitope residues recognized by each of the four antibodies ^23,26,27^ (**Ext. Data Table 1, Ext. Data Fig. 1f-g**).

**Figure 2:**
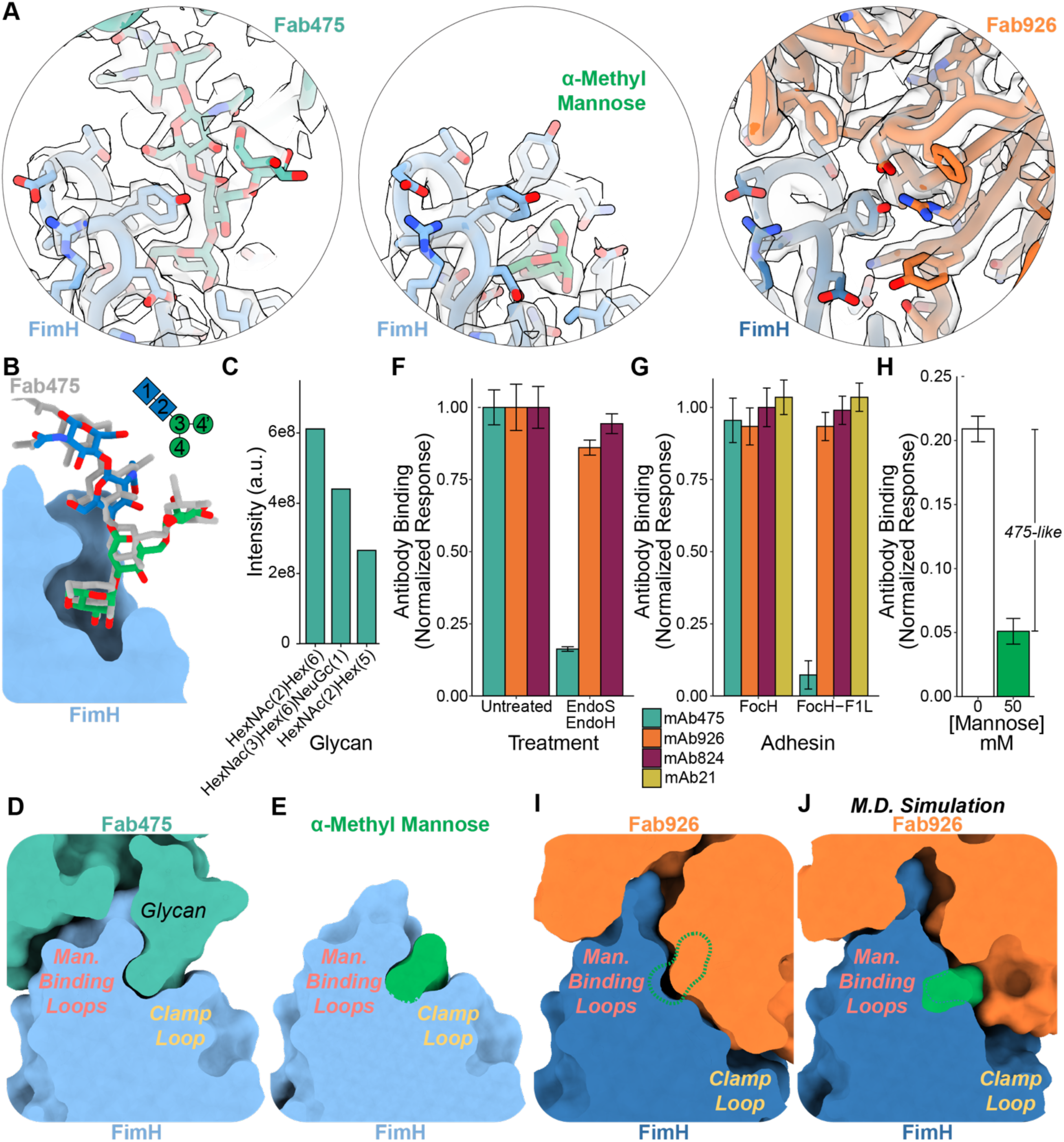
Antibodies targeting the mannose pocket prevent ligand binding via two distinct mechanisms. **A.** Volume and model of FimH ligand pockets. *Left:* FimH (light blue, sticks and ribbon) in complex with Fab475 containing and N-linked glycan (teal). *Middle:* FimH (light blue, sticks and ribbon) in complex with α-methyl mannose (green). *Right:* FimH (dark blue, sticks and ribbon) in complex with Fab926 (orange). **B**. Alignment of FimH (light blue, surface) in complex with Fab475 glycan (gray sticks) with oligomannose-3 (green/blue; PDB ID 2vco). Structures were aligned on Cα carbons 1-158 of FimH. **C**. Fab475 was separately digested using trypsin and chymotrypsin, and the resulting glyco-peptides were quantified using mass spectrometry. N-glycosylation was quantified on Asn90 of the Fab475 light chain. **D**. Slice through FimH (light blue surface) bound to Fab475 (teal surface). **E.** Slice through FimH (light blue surface) bound to α-methyl mannose (green surface). **F**. Normalized ELISA data showing changes in antibody binding to FimH caused by deglycosylation of the antibodies. N = 3 biological replicates; Error bars = combined S.D.. **G**. Normalized ELISA data comparing antibody binding to the FimH variant FocH and the mutated variant FocH-F1L. N = 3 biological replicates. Error bars = combined S.D.. **H**. Normalized and background subtracted ELISA data showing the percent mAb475-like antibodies (antibodies inhibitable by mannose). N = 3 biological replicates. Error bars = combined S.D.. **I.** Slice through FimH (dark blue surface) bound to Fab926 (orange surface) with the aligned binding location of α-methyl mannose from *E* shown as a green dotted line. **J.** Slice through the model from the molecular dynamics simulation at 50 ns; FimH (dark blue surface) bound to Fab926 (orange surface) with mannose added after simulation in green.

**Figure 3:**
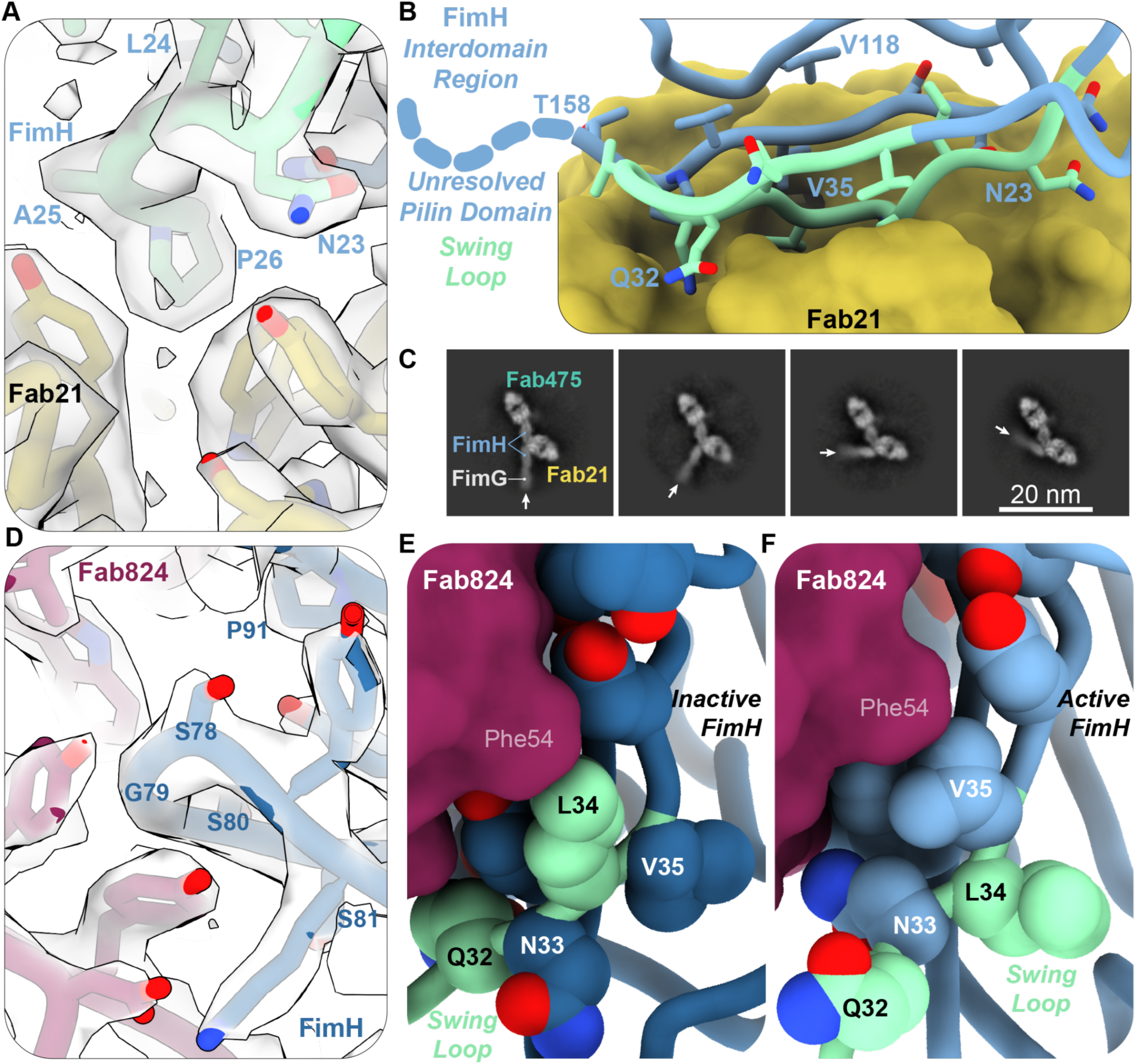
Cryo-EM structures of FimH bound to Fab21 or Fab824 provide rationale for their effects on FimH conformational dynamics. **A**. Model and volume of Fab21 (gold sticks and ribbon) binding to epitope residues on FimH (light blue/green sticks and ribbon) in the swing loop. **B**. Fab21 (gold surface) bound to FimH interdomain loops (light green sticks and ribbon, swing loop; light blue stick and ribbon, interdomain loops). Dotted light blue line represents the rest of the linker loop to the unresolved pilin domain. **C.** 2D class averages from the dataset FimH + Fab21/Fab475. Arrows point to the long axis of FimG and the FimH pilin domain. **D.** Model and volume of Fab824 (maroon sticks and ribbon) binding to core epitope residues on FimH (dark blue sticks and ribbon) in the inactive conformation. **E-F.** Fab824 (maroon surface) interacting with the FimH swing loop (green ribbon). Residues facing the antibody in the inactive conformation (**D**, light green spheres) swap positions with neighboring residues (dark blue spheres) in the active conformation (**E**, light blue spheres and ribbon).

**Figure 4:**
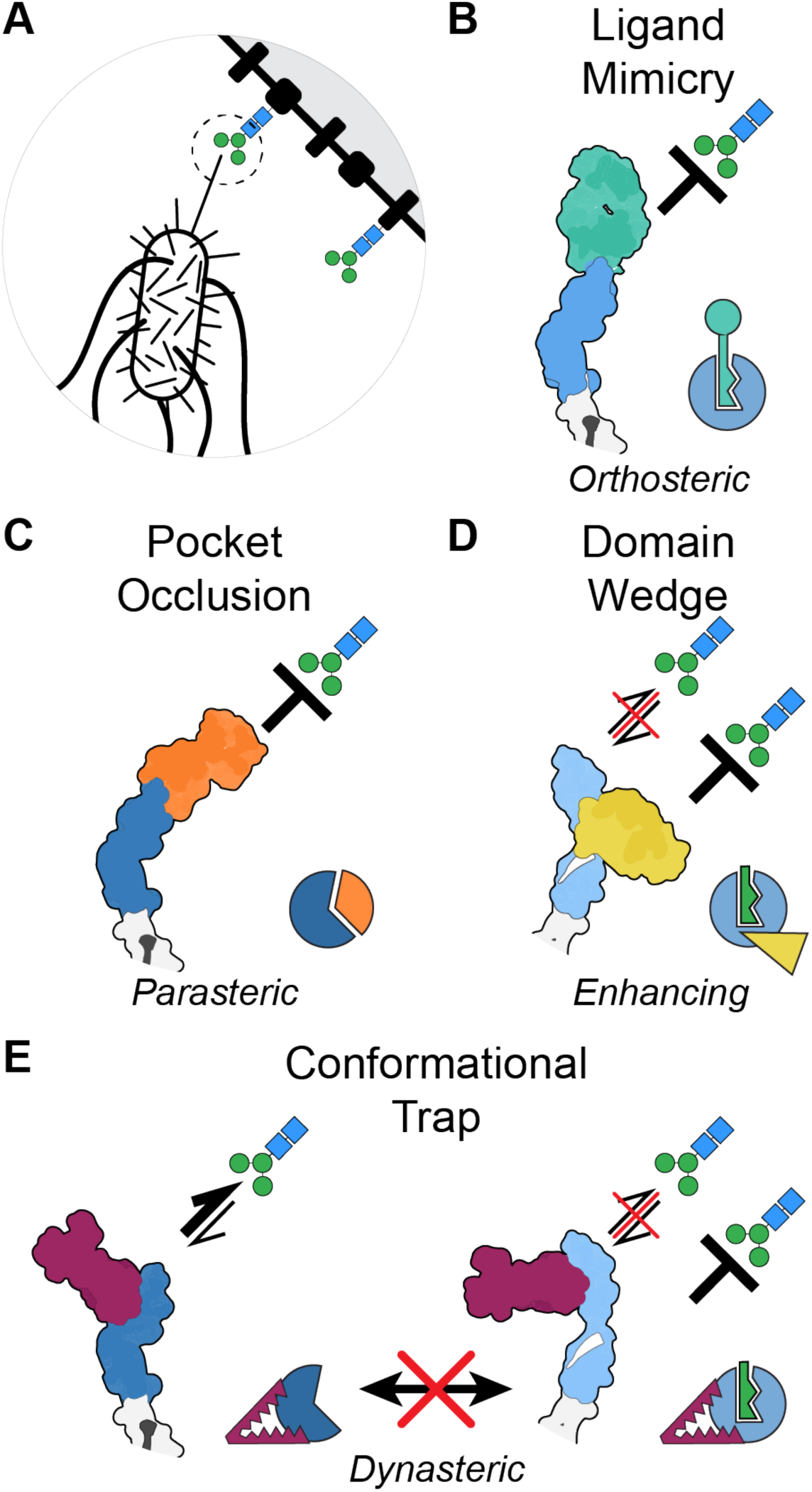
Antibodies recognizing FimH alter ligand binding with distinct mechanisms. **A**. Cartoon of bacterial fimbriae interacting with a glycosylated (blue/green) cell surface. **B**. Through a hypervariable loop glycan, Fab475 (teal) mimics FimH’s glycan ligands, blocking interactions with host glycans. **C**. Fab926 (orange) prevents glycans from binding the open pocket while also preventing pocket closure. **D**. Fab21 (gold) can only bind the active state of FimH as its epitope is inaccessible in the inactive state. Once bound, Fab21 wedges open the domains and would increase the stability of the occurring glycan-FimH interaction by stabilizing the active conformation. **E**. Fab824 (maroon) binds outside of the ligand site, but prevents conformational transitions. If trapped in the inactive state, glycans are prevented from making strong interactions with FimH. If trapped in the active state, the pocket is closed and would resist binding glycans or would increase the stability of the occurring glycan-FimH interaction.

**Table 1:**
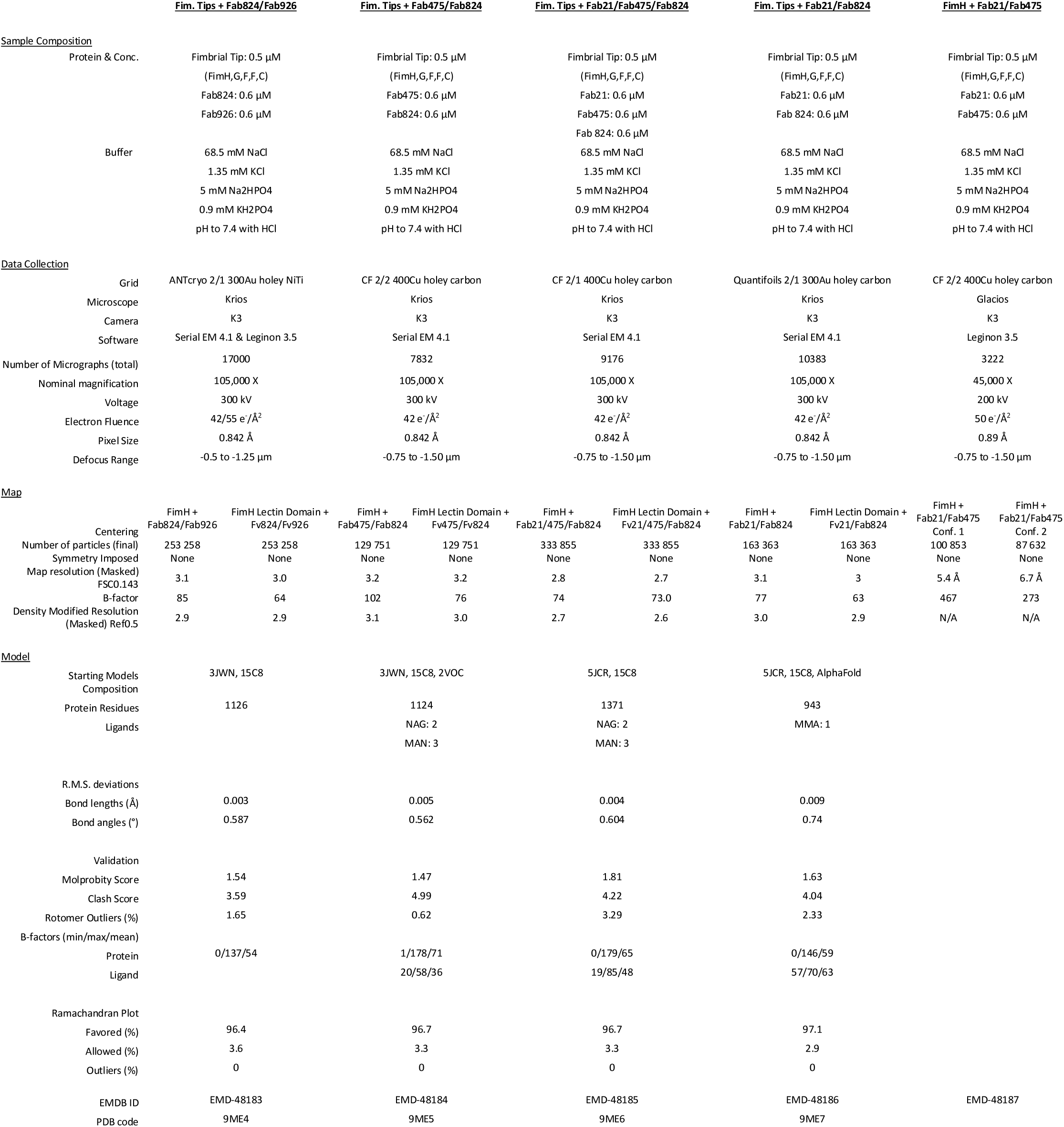
CryoEM Map and Model Statistics.

All FimH conformations observed are consistent with prior crystal structures ^7,10,12,23,26,27^ (**Figure 1e-h**). For both the FimH-Fab475/824 and FimH-Fab824/926 complexes, the lectin and pilin domains are fixed in the domain-interacting conformation at a ∼94° angle (**Figure 1e-f**). The clamp loop, which forms the external face of the ligand binding pocket, remains open in the FimH-Fab824/926 complex, and FimH adopts the inactive conformation (**Figure 1e, Ext. Data Fig. 4c**). In contrast, this loop is closed in the presence of Fab475, and the

FimH-Fab475/824 complex adopts the intermediate conformation (**Figure 1e-h**, **Ext. Data Fig. 4c)**. The lectin and pilin domains are separated and flexible relative to each other in the three complexes containing Fab21: FimH-Fab21/475; FimH-Fab21/475/824; and FimH-Fab21/824/α-methyl mannose (**Figure 1g-h, Ext. Data Fig. 5a, Movie 1**). All three complexes with Fab21 are in the active conformation.

### A glycan on mAb475 competitively inhibits ligand binding

To our surprise, the cryoEM data revealed a glycan linked to Asn90 on the Fab475 hypervariable loop 3 (complementarity determining region 3; CDR3) in the glycan-binding pocket of FimH (**Figure 2a**). The volume supports the placement of five sugars from an N-linked glycan in the structure, with the two sugars proximal to N90 likely containing N-acetyl groups and the FimH pocket containing mannose. Both the Asn90 glycan on Fab475 and the residues in the FimH ligand pocket adopt positions very similar to FimH lectin domain structures bound to oligomannose-3, recapitulating glycan and mannose binding ^6,8^ (**Figure 2a-b, Ext. Data Fig. 5b**). This result rationalizes the competitive inhibition seen between mAb475 and mannose, and how, similar to soluble mannose, mAb475 triggers the transition of FimH to the active conformation.

To identify and quantify the glycosylations on Fab475, we performed mass spectrometry on Fabs from each antibody. Only Fab475 exhibited N-glycosylation on its light chain, with two glycosylated sites present in the light chain CDRs (**Ext. Data Table 2**). The first site, Asn28 in CDR1, showed a diverse portfolio of N-linked glycans; however, over 95% of the glycan intensity consisted of sialylated glycans (**Ext. Data Fig. 6a**).

Inspection of the cryoEM data surrounding Asn28 showed additional disordered volume in agreement with the mass spectrometry data (**Ext. Data Fig. 5c**). The second site, Asn90 in CDR3, identified only three glycans attached to the asparagine (**Figure 2c**). Two of these are high-mannose glycans, and the third is a sialylated glycan. Investigation of deamidated asparagines generated by PNGase F treatment corroborates these identified glycans and the lack of N-glycosylation on the other Fabs (**Ext. Data Table 3**). Further confirmation of glycosylation on Fab475 and the absence of glycosylation on the other Fabs was obtained through native mass spectrometry analysis. Intact analysis of Fab475 revealed highly heterogeneous spectra that eluded simple deconvolution, which is characteristic of glycosylated proteins (**Ext. Data Fig. 6b**). In contrast, Fab926, Fab824, and Fab21 revealed less complex and interpretable spectra with no indication of glycosylation.

While the glycan only accounts for half of the buried surface area, it dominates the mAb475-FimH interaction. In total, Fab475 buries 712 Å^2^ of the FimH pocket (**Ext. Data Fig. 4d, Ext. Data Table 1**). The glycan on Fab475 buries 372 Å^2^ of surface area, which is nearly identical to the buried area of a bound glycan alone (PDB ID 2vco, 379 Å^2^) ^6^. Dissociation constants for oligomannose-3 and oligomannose-5 are 10-20 nM, suggesting that the boost in affinity observed for Fab475 (4 nM) arises from additional interactions with the antibody ^6,26^. The remainder of the interactions occur between the hypervariable loops in the heavy chain of Fab475 and FimH. These contacts are primarily made with the outer side and top of the mannose binding loop 136-140 through a series of polar interactions (**Ext. Data Fig. 5d**). Using both its covalently attached glycan and heavy chain residues, Fab475 grabs the mannose binding loops of FimH in the active conformation like pincers (**Figure 2d-e**).

We assessed the importance of the glycan in mAb475 binding and inhibition of FimH by deglycosylating mAb475 and performing binding experiments. Unlike the light chains of the other three antibodies, the light chain of mAb475 migrated at approximately 30 kDa on SDS-PAGE, 5 kDa heavier than the predicted 25 kDa (**Ext. Data Fig. 7a, Supp Data 1**). Treatment of mAb475 with endoglycosidase S (Endo S), which cleaves off sialylated N-linked glycans, shifted the light chain to a lower molecular weight, suggesting complete removal of the targeted glycan. However, removal of this glycan did not affect mAb475 binding to FimH (**Ext. Data Fig. 7a, Supp Data 1**). Treatment of mAb475 with endoglycosidase H (Endo H), which cleaves off high-mannose and some hybrid glycans, shifted a portion of the light chain to a lower molecular weight, indicating partial removal of the targeted glycan. This partial deglycosylation decreased mAb475 binding to FimH (**Ext. Data Fig. 7a, Supp Data 1**). Treatment with both Endo S and Endo H fully shifted the molecular weight of the light chain closer to the predicted size of 25 kDa (**Ext. Data Fig. 7a, Supp Data 1**) and eliminated mAb475’s ability to bind FimH (**Figure 2f, Ext. Data Fig. 7b**). Along with the mass spectrometry and cryo-EM results, the biochemical results confirm that the complex glycans primarily found on Asn28 do not contribute to FimH binding while the high-mannose glycans on Asn90 drives binding and inhibition by mAb475.

### mAb475-like antibodies are naturally elicited

The FimH-Fab475 interaction presents a previously unobserved antigen-antibody interface where a hypervariable-loop glycan is specifically recognized by the target lectin, allowing the antibody to directly block the ligand-binding site. However, mAb475 is not unique; we previously identified six additional antibodies with competitive, orthosteric binding behavior similar to that of mAb475 ^23^. Sequence alignments of these antibodies and closely related sequences from public databases show that the antibodies from our study contain a glycosylation motif (NxS/T, x = all amino acids except proline) at the aligned equivalent residue of Asn90 (**Ext. Data Fig. 7b**). In addition, the sequences segregate into two groups with the second glycosylation site at Asn28 appearing in one group but not in the second. Together, this suggests that the mAb475-like antibodies in our study originated from two independent germline lineages and further affinity maturation produced multiple glycosylated antibodies.

The approximate percentage of mAb475-like antibodies in the polyclonal antibody pool of mouse antiserum obtained by immunization with the FimH lectin domain was estimated as follows. We selectively depleted antibodies from aliquots of antiserum using bacteria expressing: a) fimbria without FimH; b) fimbria with the FimH-FocH, a variant that primarily adopts the active conformation; and c) fimbria with the FimH-FocH^F1L^ mutant, where the phenylalanine at position 1 is replaced with leucine, eliminating the ability of FimH to bind mannose and glycans. None of the monoclonal antibodies recognized fimbriae lacking FimH. All four antibodies recognized FimH-FocH, but mAb475-like antibodies do not recognize FimH-FocH^F1L^ (**Figure 2g, Ext. Data Fig. 7c**).

After adsorption, we probed for FimH binding within the remaining pool of antibodies in each antiserum sample. Adsorption with bacteria lacking FimH did not significantly decrease anti-FimH antibodies in the serum (**Ext. Data Fig. 7d**). In addition, mannose had no significant effect on unadsorbed antiserum or antiserum treated with bacteria lacking FimH (**Ext. Data Fig. 7d**). In contrast, the vast majority of the antibody pool was removed by bacteria expressing FimH-FocH fimbriae (**Ext. Data Fig. 7d**). A small but measurable increase in the signal was observed in the presence of mannose, indicating that some of the non-absorbed antibodies may be activating, like mAb21 (**Ext. Data Fig. 7d**). When the antiserum was absorbed with bacteria expressing the FimH-FocH^F1L^ mutant, a larger portion of the antibodies remained unabsorbed (**Figure 2h, Ext. Data Fig. 7d**). Notably, the binding significantly decreased in the presence of mannose, indicating that the bulk of the non-adsorbed antibodies compete with mannose for binding to FimH. This suggests the polyclonal antibody response contains a minor but clearly detectable portion of orthosteric, mAb475-like antibodies (**Figure 2h**).

### mAb926 sterically blocks ligand pocket closure

In contrast to mAb475 binding within the closed ligand pocket, mAb926 binds the open ligand pocket with FimH in the inactive conformation (**Figure 1a,e** and **2a, Ext. Data Figure 4b-c**). The binding epitope of Fab926 centers around the mannose-binding pocket in the open conformation, with a total interaction area of 983 Å^2^ (**Ext. Data Fig. 4d, Ext. Data Table 1**). Both chains of Fab926 form polar interactions and hydrophobic packing interactions with the inner and outer faces of the mannose binding loops of FimH (**Figure 2a, Ext. Data Fig. 8a**). This configuration positions Fab926 to bury amino acids from portions of hypervariable loops on the heavy chain in the open ligand binding pocket. Additional sections of the paratope sterically block clamp loop closure without making specific contacts with the clamp loop (**Data Fig. 8b-c**). Together, the heavy and light chains of Fab926 pinch the mannose binding loops of the ligand pocket while the heavy chain fills and splays open the binding pocket (**Figure 2i**). The differences in FimH’s conformation and binding epitopes between mAb926 versus mAb475 binding explain why the antibodies allosterically stabilize inactive and active states of FimH, respectively.

mAb926 binding shows some similarities to both glycan and mAb475 binding. As with Fab475, hypervariable loops on Fab926 contact the top and the outer side of the mannose binding loops. The area buried in the ligand pocket by Fab926 overlaps with regions previously shown to be buried by mannose, high-mannose glycans, and small-molecule mannosyl compounds ^6,18,29,30^ (**Ext. Data Fig. 8d-e**). Hypervariable loops on Fab926 recreate a subset of the polar interactions generated by glycan ligands, including contacts made by the mannose moiety deep in the binding pocket and others made by the N-acetylglucosamines with the upper portions of the mannose binding loops (**Ext. Data Fig. 8e**). In addition, several hydrophobic residues on Fab926 contact a hydrophobic patch in the FimH pocket (**Ext. Data Fig. 8f**). In sum, the structure demonstrates that fully bound Fab926 sterically occludes native ligand binding, and the closure of the binding pocket required for the active state.

### The mAb926 - FimH interface is flexible

While the cryoEM structures show how mAb475 and mAb926 stabilize different FimH conformations, the models do not explain why mAb926 inhibits FimH in a non-competitive manner. Previous work demonstrated that mannose is about 30 times less effective at blocking mAb926 binding as compared to mAb475 binding, even though both antibodies’ epitopes include the ligand-binding site ^26^ (**Figure 1a-b**). In addition, mAb926 and mannose can reciprocally displace each other from FimH, which is not observed with mAb475 ^26^. These previous results suggest more flexibility at the FimH-Fab926 interface than is represented in our single structure.

To explore the flexibility of Fab926 binding, we used molecular dynamics simulations. We performed simulations using either the FimH-Fab926 structure or the FimH-Fab475 structure to determine the range of flexibility for each interaction. Over the course of a 50-ns simulation run at 330K, the FimH-Fab475 interaction remained unperturbed, and the clamp loop remained closed (**Ext. Data Fig. 9a-b**). In contrast, the simulation with FimH-Fab926 showed disruption of interactions between the FimH lectin binding pocket and hypervariable loop 1 of Fab926 (residues 26-33), which sits in the lectin domain pocket and above the clamp loop (**Figure 2i, Ext. Data Fig. 9c-e**). The portions of this loop outside of the binding pocket, amino acids 24-28, are not well resolved in the cryoEM structure, also suggesting flexibility (**Ext. Data Fig. 8b-c**). Coincident with the movement of the Fab926 loop, the clamp loop of FimH moves towards the closed position adopted in the active conformation, in the direction of Fab926. Together, these conformational changes would create space for mannose in the binding pocket while mAb926 is still bound to FimH, and the simulations highlight flexibility within the FimH-mAb926 binding interface (**Figure 2j**). The simulations suggest that mannose may bind FimH while Fab926 is bound, and Fab926 may bind when mannose is bound, with each destabilizing the binding of the other.

### FimH retains interdomain flexibility when bound to mAb21

The structures containing Fab21 support previous work showing mAb21 binds FimH in the active conformation at functional epitope residues located in the interdomain interface ^23,31^ (**Ext. Data Fig. 5a, Ext. Data Table 1**). The 721 Å^2^ Fab21 epitope only includes residues on the interdomain interface of the lectin domain (**Figure 3a-b, Ext. Data Table 1**). The antibody binds in between the FimH lectin and pilin domains but does not specifically interact with the pilin domain (**Ext. Data Fig. 5a**). The two chains of Fab21 pinch swing loop residues 23-29 and pack tightly against the remainder of the swing loop and the interdomain loops (**Figure 3a-b**). In the high-resolution structures containing Fab21, Fab824, and either Fab475 or α-methyl mannose, FimH is in the active conformation and Fab21’s position physically blocks the pilin and lectin domains from reengaging and forming the fixed interface seen in the intermediate and inactive conformations (**Ext. Data Fig. 5f**).

The pilin domain remains flexible when Fab21 is bound. The class averages containing Fab21 and Fab475 show the pilin domain in multiple positions (**Figure 3c, Movie 1, Ext. Data Figure 3e** and **5a**). Reconstructing the high-resolution datasets containing Fab21 without masks and filtering the volumes only resolves the portion of the pilin domain proximal to the lectin domain, demonstrating that it is present but blurred in the datasets, a signature of mobile domains (**Ext. Data Fig. 5e**). Fab21 may block a subset of possible lectin-pilin orientations, but the lectin domain retains its ability to flex and rotate relative to the pilin domain.

### mAb824 traps dynamic residues on FimH

As shown previously, the Fab824 epitope spans the surface of the beta sheet distal to the ligand binding pocket of the FimH lectin domain (**Ext. Data Fig. 1g, Ext. Data Table 1**). The light and heavy chains of Fab824 straddle residues 79-82 and 91, originally identified as critical interactions in a mutational epitope screen ^27^ (**Figure 3d, Ext. Data Fig. 1h**). Additionally, the full contact surface of Fab824 bound to FimH contains residues that were previously identified as part of the epitope by ^13^CH_3_-NMR ^27^ (**Ext. Data Fig. 1g**). The epitope defined by the contact surface in the cryo-EM structure includes 983 Å^2^ of the FimH surface in the inactive conformation (**Ext. Data Fig. 4d, Ext. Data Table 1**). This contact surface is larger than the epitope of Fab824 in the active conformation (854 Å^2^, previously defined by NMR; 839 Å^2^, this study) ^27^ (**Ext. Data Fig.** 4d, Ext. Data Table 1).

The models provide a structural rationale for the previous observation that mAb824 can bind all conformations of FimH but prevents the conformational transition between active and inactive conformations ^27^. mAb824 directly interacts with residues on FimH that flip between the surface and core of the protein during conformational transitions (**Ext. Data Fig. 1g**). FimH residues Leu34 and Val35 have been identified as a “toggle switch” in which the two side chains flip around the beta strand axis, essentially swapping places with each other in the two conformational states [2021KisielaMagalaetalPLoSPath]. In the inactive FimH/Fab824 structure, Phe54 from Fab824 packs against the side chain of Leu34 while Val35 points into the core of FimH (**Figure 3e**). In the active state, the side chain of Val35 is on the surface of FimH and packs against Phe54 from Fab824 (**Figure 3f**). Fab824 also makes contacts with the side chains of Tyr108 and Lys76 on the adjacent strands within the same beta sheet (**Ext. Data Fig. 8g**). Three epitope residues on separate strands of the beta sheet, Val36, Leu107, and Val75, do not interact with Fab824 but pack tightly against each other. Together, the packing of mAb824 and the beta strand residues pin the beta strands in place, preventing the formation of the 6-Å gap required for the Leu34/Val35 switch. Thus, by interacting with a large set of dynamic residues and packing directly against Leu34 in the inactive state or against Val35 in the active state, Fab824 prevents the rearrangements required for transition between the two conformations.

## Discussion

The high-resolution structures of FimH bound to antibody fragments provide a structural framework to interpret prior results in the context of a bacterial infection (**Figure 4a**). The results demonstrate four distinct effector mechanisms by which antibodies perturb FimH. Orthosteric mAb475 directly mimics native FimH ligands by placing a glycan deep into the closed pocket of FimH in the active conformation (**Figure 4b**). In contrast, parasteric mAb926 occludes the ligand pocket by interacting broadly within and near the open pocket of inactive FimH (**Figure 4c**). The attachment-enhancing mAb21 can only bind FimH in the active conformation and prevent FimH from transitioning back to the inactive conformation, trapping the bound ligand (**Figure 4d**). Dynasteric mAb824, acting as a conformational trap, holds FimH in either the high or low-affinity conformation by contacting key allosteric residues (**Figure 4e**). These insights into the mechanisms of neutralization of cell attachment proteins will likely facilitate future development of next-generation therapies.

mAb475 presents a previously unreported, yet likely common, mechanism of interaction between antibodies and lectin domains via a “functional glycan” (**Figure 2a** and **d**). Hypervariable loop glycosylations occur in 15% of circulating immunoglobulin G antibodies, and while several hypotheses have been proposed to explain why they exist, none included the direct, specific engagement of the glycans with the target antigen ^32–35^. Our data suggest that mAb475-like antibodies are a general response strategy against invaders and may partly explain the substantial percentage of hypervariable loop glycosylations. Many human pathogens, including bacteria, fungi, and eukaryotic parasites, attach to their host using oligosaccharide-binding lectin-like domains ^3^. During infection, lectins likely trigger the development of antibodies with glycosylated hypervariable loops. These antibodies block microbe binding by competitively inhibiting the binding of glycans on epithelial cell surfaces. We believe that the interaction between mAb475 and FimH provide a starting point for the development of immune therapies targeting adhesins or other oligosaccharide-binding proteins.

Glycosylated hypervariable loops on surface immunoglobulins are also a hallmark of several diseases, including a subset of follicular lymphomas ^36–38^. Diffuse large B-cell lymphomas present with B cell receptors containing N-linked, high mannose glycans in the hypervariable loops, but how these glycosylation sites arise in the first place remains a subject of debate ^34,38,39^. Comparison of variable region fragments from a B cell receptor from a diffuse large B-cell lymphocyte show a high mannose glycan at the equivalent residue of Asn90 as well as other paratope locations ^38,40^ (**Ext. Data Fig. 5g**). Our work suggests that a polyclonal antibody response to an infection involving lectin-like adhesins would select for hypervariable loops containing high mannose glycans, creating a pool of B cells with this type of glycan. Further mutations and the breaking of tolerance would generate cells capable of transforming into cancer ^39^. More broadly, microbial selection of hypervariable loop glycans could play a role in autoimmune diseases that have been associated with antibody glycosylation, including rheumatoid arthritis and primary Sjögren’s syndrome ^41,42^. Thus, bacterial selection of glycosylated hypervariable loops during infection may represent one step on the pathway to a subset of cancers.

The FimH-Fab475 and FimH-Fab926 structures provide a clear rationale for differences in the allosteric effects of these two categories of antibodies; however, the FimH-Fab926 model does not explain the unique and unusual properties of parasteric antibodies that imply their possible, transient binding to FimH simultaneously with the ligand ^26^. Molecular dynamic simulations provide some clues. The simulations predict flexibility in the mAb926 hypervariable loops, with changes in loop position large enough to allow enough space for mannose to bind while mAb926 is still bound. Additional simulations and rigid body docking suggest one mechanism by which mAb926 could push ligands out of the FimH pocket. A previously published simulation demonstrated that the clamp loop of FimH serves as a molecular latch for the main mannose-interacting loops ^43^. When closed, the clamp loop stabilizes the closed pocket conformation and, thus, ligand binding; when open, ligands can leave the binding pocket. When we used models of FimH from this simulation with the clamp loop opened or closed and rigid-body dock Fab926 to check for fit, the resulting models suggested that the opening of the clamp loop creates the necessary binding space for Fab926, with fewer clashes observed between Fab926 and FimH in the open-clamp-loop conformation (**Ext. Data Fig. 9f-g**). Taken together, mAb926 may bind FimH when the FimH clamp loop samples the open conformation with mannose still present. Binding of mAb926 would prevent the clamp loop from closing and would destabilize mannose binding.

Parasteric mAb926 may provide a widely applicable blueprint for the rational design of inhibitors targeting the open binding pocket. mAb926-like antibodies are likely widespread; a recent unpublished study identified an antibody binding the open pocket of FimH ^44^. Additionally, regions resembling the clamp loop can be identified in distant homologues of FimH, including the FimH of *Klebsiella pneumoniae* and the GalNAc-specific adhesin, FmlH, in F9 fimbriae of *E. coli*. A similar “dock, lock, and latch” mechanism of ligand binding occurs in other cell attachment proteins and the latch elements have been shown to be good protective vaccine targets ^45–48^. Our structure with Fab926 also shows that targeting the open pocket of FimH is a viable strategy to block glycan binding without the structural rearrangements required for the active conformation (**Figure 2a** and **h**). The stable active conformation of FimH requires continuous separation of the lectin and pilin domains, which is difficult to achieve without the application of tensile force ^7^. Keeping the pocket open also provides a larger antibody target; the Fab926 and Fab475 structures suggest that fitting a hypervariable loop into the closed ligand binding pocket of FimH would be difficult due to the space constraints. Targeting the open pocket and the clamp loop of adhesins with antibodies, synthetic peptides, or small-molecule compounds to destabilize or prevent ligand binding represents an effective way to block and reverse pathogen adhesion.

Our study confirms the long-standing hypothesis that FimH-activating mAb21 acts as a wedge between the lectin and pilin domains by binding loops on the lectin domain within the domain interface in the active conformation ^23,31^. Fab21 specifically binds to the active conformation of the dynamic interdomain and swing loops on the lectin domain (**Figure 3b**). By binding in this location, mAb21 allosterically stabilizes the active conformation and sustains ligand binding by FimH without blocking the interdomain flexibility of the lectin and pilin domains ^31^.

The FimH-Fab824 structure provides insight into the dynasteric mechanism of action. As previously predicted, the structures show that mAb824 binds very similar epitopes of FimH in the active, intermediate, and inactive conformations ^27^. This conformational blindness combined with Fab824’s unmeasurably low dissociation rate results in Fab824 trapping FimH in whichever conformation it first encounters ^27^ (**Figure 3e-f**). The structures confirm that the antibody epitope on FimH includes dynamic residues that undergo characteristic and essential conformational changes during the transformation between the inactive and active conformations. The tight mAb824-FimH interactions within this region likely impose severe steric restrictions on the ability of FimH to undergo the allosteric switch, which likely involves transient beta strand separation. mAb824 traps either the active or inactive conformation of FimH, meaning its effects on binding are mixed. Although the effect of mAb824 in an infection model *in vivo* has not yet been demonstrated, the conformational trapping would essentially paralyze the adhesive function of FimH that involves dynamic switching between the inactive and active states.

In sum, we present high-resolution structures of FimH in complex with Fab fragments which shed light on multiple distinct mechanisms of antibody-antigen interactions and can serve as blueprints for conceptually novel anti-adhesion therapies. Along with previous work, this study shows how a set of monoclonal antibodies derived from a pool of polyclonal antibodies recognize different adhesin epitopes and bind simultaneously yet have distinct consequences for the behavior of an allosterically driven process. Together, the antibodies can produce either additive or opposing effects on adhesion. Paradoxically, the outcomes generated by the antibodies may not always be beneficial to the host and must therefore be considered in the context of antigens used in vaccine development.

## Acknowledgements

We thank Dagmara Kisiela, a former member of Sokurenko lab, for developing multiple protocols and purifying key reagents used in this study, as well as the overall intellectual impact into the concepts of orthosteric, parasteric and dynasteric antibodies. We thank Angelo Ramps for technical support with sample preparation. We thank the Arnold and Mabel Beckman Cryo-EM Center at the University of Washington for electron microscope use. We thank Joel Allen for helpful discussions on antibody glycosylations and members of the Kollman group for valuable feedback provided during cryo-EM data collection, processing, and data presentation. This work was supported by the US National Institutes of Health (grant nos. F32 AI145111 to K.L.H., K99 GM141364 to P.M., R00 GM147304 to N.M.R., R01 AI171570 to E.V.S. and R.E.K., R35 GM149542 and S10 OD023476 to J.M.K.). The molecular dynamics simulations were performed on the Expanse supercomputer at the San Diego Supercomputing Center thanks to a ACCESS allocation with grant number TG-BIO240017 (to G.I.).

## Competing Interests Statement

EVS and VT hold US10722580B2 patent ‘Compositions and methods for treatment and prevention of uropathogenic E. coli infection’ that include therapeutic use of mAb475 and mAb926. N.M.R. receives support from Thermo Fisher Scientific under a nondisclosure agreement and is a consultant for Tegmine Therapeutics, Cartography Biosciences, and Augment Biologics.

## Data Availability Statement

The cryo electron microscopy maps generated for this manuscript are available from the EMDB (https://www.ebi.ac.uk/emdb/) at the accession codes listed in Table 1 of the manuscript (EMDB IDs: EMD-XYZ1, EMD-XYZ2, …). The protein models generated for this manuscript are available from the RCSB PDB (https://www.rcsb.org/) at the accession codes listed in Table 1 of the manuscript (PDB IDs: XYZ1, XYZ2). The mass spectrometry proteomics data have been deposited to the ProteomeXchange Consortium via the PRIDE partner repository with the dataset identifier PXD057683 (Username; Password: NcMsYBw7jsAA).

## Methods

### Expression and Purification of Fimbrial Tips from E. coli

The genes encoding for FimC (with a his-tag at the c-terminus) and FimF-FimG-FimH (FimH UniProt ID: P08191) of the fimbrial tips were cloned into pRSET-B expression plasmid as previously described ^13^. The plasmid was transformed into BL21 (DE3) cells via electroporation and plated on LB agar overnight. The cells were grown at 37°C in 1 L cultures until the optical density at 600 nm (OD600) reached 0.5-0.7. Protein expression was induced by the addition of 0.5 mM IPTG, and the cultures were grown at 22°C overnight. Cells were harvested by centrifugation and resuspended in PBS buffer containing 5 mM imidazole. The cells were lysed using a French press at 1000 psi, and the lysate was clarified by centrifugation. The clarified lysate was loaded onto a nickel column and washed with PBS buffer containing 5 mM imidazole. A second wash step was performed with PBS buffer containing 50 mM imidazole. The fimbrial tips were eluted with PBS buffer containing 100 mM imidazole. The protein was then further purified by size exclusion chromatography on a Superdex 75 column. Fractions containing the fimbrial tips were pooled and concentrated.

### Generation of Monoclonal Antibodies and Fab Fragments

The lectin domain of FimH was used to raise the monoclonal antibodies as previously reported ^49^. The mAbs were expressed and purified from hybridoma cell cultures. The cell cultures were clarified by centrifugation, and the supernatant was loaded onto a Protein G-agarose column. The column was washed with IgG binding buffer, and the mAbs were eluted with IgG elution buffer from Thermo Fisher Scientific. The mAbs were further purified by size exclusion chromatography on a Superdex 200 column in PBS buffer. Purified mAbs were then digested to Fab fragments using agarose-immobilized ficin from Thermo Fisher Scientific and purified by size exclusion on a Superdex 75 column.

### Fimbrial Tip-Fab Complex Assembly

The fimbrial tips/Fab complexes were generated from a mixture of fimbrial tips and Fab fragments in a 1:1.2 ratio. All protein concentrations were measured at UV 280 nm using a NanoDrop.

### Cryo-Electron Microscopy

Protein samples were assembled at the ratio listed above and at the concentrations listed in Table 1. Protein solutions were applied to glow-discharged grids (CF - Protochips Inc.; ANTcryo – Single Particle, LLC; Quantifoil, SPT Labtech), blotted, and plunge-frozen in liquid ethane using a manual plunging apparatus at room temperature. Data collection was performed using an FEI Titan Krios transmission electron microscope operating at 300 kV (equipped with a Gatan image filter (GIF) and post-GIF Gatan K3 Summit direct electron detector) or an FEI Glacios (equipped with a Gatan K3 Summit direct electron detector) using the software packages Leginon (v3.5) or SerialEM (v4.1) [2005SulowayetalJStructBiol, MastronardeXYZ].

### Data Processing

Movies were aligned, corrected for beam-induced motion, dose-weighted, and binned (2x – if collected in super-resolution mode) using Patch Motion Correction in cryoSPARC (v4.4.1) or cryoSPARC Live^50^. CTF parameters were estimated using Patch CTF ^50^. For the datasets FimH + Fab824/926, FimH + Fab21/824, and FimH + Fab21/475/824, micrographs were excluded if they fell outside of these thresholds: −0.1 to −2.5 nm defocus; a CTF fit of 2-6; and a relative ice thickness of 1.050-1.250. For datasets FimH + Fab475/824 and FimH + Fab824/926, particles were picked using Blob Picker with these parameters: 100-300 Å min-max diameter with a separation distance of 0.4 diameters using an elliptical blob. For datasets FimH + Fab21/824 and FimH + Fab21/475/824, particles were template picked with templates generated from a low-resolution volume of Fab21/475/824. Binned particles (4x) and two rounds of 2D Classification were used to remove noise and poorly aligning particles, and the remaining particles were used in *ab initio* Reconstruction and 3D Homogenous Refinement. Particles were re-extracted with decreased binning (1.3x), and additional rounds of 3D refinement were completed, including Homogenous Refinement, Non-Uniform Refinement, and Local Refinement. Datasets were sent through a round of 3D Classification with hard classification and masks as shown in Ext. Data Fig. 2, followed by additional 3D refinement with an FSC cutoff of 0.143. The final refined, unsharpened half maps were exported to Phenix (v1.20) where density modification and additional resolution estimation was performed (FSC cutoff of 0.5) ^51,52^. Local Resolution Estimation was performed in cryoSPARC using the maps generated in the final round of refinement and mapped onto the density modified volume in ChimeraX (v1.7) ^53^.

### Model Building

PDB ID 3JWN and 15C8 were used as initial models for FimH and Fab475, 824, and 926 ^13,54^. The correct FimH sequence was manually added to 3JWN and Fab sequences were threaded onto 15C8 using the Modeller function in ChimeraX ^55^. AlphaFold 3 was used to generate an initial model of Fab21 ^56^. Models were iteratively refined using a combination of ISOLDE (v1.7) in ChimeraX (v1.7), Coot (v0.9.8.7), and phenix.real_space_refine and phenix.validation_cryoem in Phenix ^52,53,57–59^. The phenix.validation_cryoem tool was used to generate model statistics.

### Image Generation and Calculations for Micrographs and Protein Models

Cryo electron micrographs were motion corrected using Patch Motion Correction in cryoSPARC or cryoSPARC Live (v4.4.1) and Gaussian filtered and contrast adjusted in Fiji (2.14.0/1.54f). ChimeraX (v1.7) was used for: protein volume and model visualization, volume alignment (Fit to Model tool or the “fit” command), model alignment (Matchmaker tool or the “align” command), buried surface area calculations (“measure buriedarea” command), hydrophobicity calculations (“hydrophobic coloring” tool), and hydrogen bonds identification (“hbond” tool). All electron micrograph image, volume, and protein model figures were assembled in Adobe Illustrator CC6 (v26.0.1).

### Glycopeptide sample processing

Each of the Fab molecules was prepared separately in parallel. Each aliquot of individual Fabs contained 5 µg of the Fab molecule in 200 mM Tris buffer solution, was subjected to a 5-minute heat shock at 95°C, was reduced with 5 mM tris(2-carboxyethyl)phosphine hydrochloride (TCEP) for 30 minutes at 60°C, and was alkylated with 20 mM chloroacetamide, followed by a 15-minute quench using TCEP. Sodium deoxycholate was added to a final concentration of 1%. Proteins were digested overnight at 37°C with either trypsin (1:50 μg/μg protease:protein) (Promega) or Chymotrypsin (1:70 µg/µg protease:protein) (Promega). The SDC in all samples was precipitated twice with 2% formic acid (FA), after which samples were desalted using Strata-X cartridges (Phenomenex) by conditioning the cartridge with 1 mL acetonitrile (ACN) followed by 1 mL 0.2% formic acid (FA) in water. Peptides were then loaded onto the cartridge, followed by a 1 mL wash with 0.2% FA in water. Peptides were eluted with 400 μL of 0.2% FA in 80% ACN and dried via vacuum centrifugation. Individual trypsin or chymotrypsin digestions were repeated for each Fab in parallel. These samples were further processed using a PNGase F treatment, where samples were heat-shocked for 5 minutes at 95°C to inactivate proteases before they were incubated with 500 units of PNGase F (New England Biolabs) for 4 hours at 37°C. Following PNGase F treatment, these samples were desalted using the same protocol as above and dried via vacuum centrifugation. After drying, all samples were stored at −70°C until mass spectrometry analysis.

### Glycopeptide data acquisition

Glycopeptides were analyzed on an Orbitrap Ascend Tribrid Mass Spectrometer (Thermo Fisher Scientific) coupled to a Vanquish Neo UHPLC (Thermo Fisher Scientific). First, samples were reconstituted in 0.1% FA and loaded on a C18 trap column (300 µm x 5 mm, 5 µm particles, PepMap Neo) using a 300 nL/min flow using buffer A (0.1% formic acid). Subsequently, samples were linearly eluted using a 55-minute gradient ranging from 2.2% buffer B (99.9% acetonitrile with 0.1% FA) to 28% buffer B on a C18 analytical column (75 µm x 25 cm, 1.7 µm particles, IonOptics). The column was washed for 5 minutes with 99% B after separation and equilibrated with 100% buffer A. Precursors were ionized using a nanospray flex ionization source (Thermo Fisher Scientific) held at +2.0 kV compared to ground, and the inlet capillary temperature was held at 275°C. The instrument method collected spectra in a data-dependent fashion using full scan (MS1) settings of 120,000 resolving power at 200 m/z, a normalized AGC target of 250% (1,000,000 charges), and an MS1 mass range from 400-1800 m/z. Dynamic exclusion was set at 15 seconds, and only charges 2-6 were selected for fragmentation. Precursor ions were isolated for MS/MS scans using a 0.7 m/z quadrupole isolation width, measured in the Orbitrap with an MS2 resolution of 30,000 at 200 m/z, a normalized AGC target set to 200% (100,000 charges), and a maximum injection time of 59 ms. Selected precursor ions were fragmented with stepped collision energy/higher-energy collisional dissociation (sceHCD) performed using 20, 30, and 40% normalized collision energies. sceHCD scans containing at least four glycan-specific oxonium ions (126.055, 138.0549, 144.0655, 168.0654, 186.076, 204.0865, 274.0921, 292.1027, and 366.1395 m/z) triggered electron-transfer/higher-energy collision dissociation (EThcD) scans, where precursor ion isolation was performed with the quadrupole using a 1.6 m/z isolation window, an AGC target of 200% (100,000 charges), and a maximum injection time of 251 ms. EThcD fragmentation was performed using a 50 ms reaction time and a 2e6 reagent target.

### PNGase F-treated peptide data acquisition

The PNGase F-treated glycopeptides were analyzed using the same LC-MS setup and gradient as the glycopeptides. The instrument method used an MS1 resolution of 120k at 200 m/z, a normalized AGC target of 100% (400,000 charges), and an MS1 mass range ranging from 375-1600 m/z. Dynamic exclusion was set at 30 seconds, and only charges 2-6 were selected for fragmentation.

HCD MS/MS scans were performed using a 0.7 m/z quadrupole isolation width, and fragment ions were mass-analyzed in the Orbitrap with an MS2 resolution of 15k at 200 m/z. HCD MS/MS scans used a normalized AGC target set to 100% (50,000 charges), a maximum injection time of 27 ms, and a normalized collisional energy of 30%.

### Glycopeptide data analysis

Raw data were searched using Byonic ^60^. Each glycopeptide search for N-glycosylation was conducted separately, each using the fasta sequence file specific to the Fab. The N-glycan database consisted of 309 unique mammalian N-glycan compositions. Carbamidomethylation was set as a fixed modification, and oxidation on M, deamidation on N, and acetylation on the N-term were used as variable modifications. A maximum of 3 miscleavages were allowed, and a maximum of three glycosites were allowed for any one glycopeptide. Glycan quantification was performed on the MS1 level using Skyline ^61^. Data was graphed using R (version 4.4.0). Filtering metrics included a Byonic score greater than or equal to 200, and a logProb value greater than or equal to 2.

O-Pair Search was performed in MetaMorpheus ^62,63^. The ‘Glyco Search’ option was selected, and the O-glycopeptide search feature and the Oglycan were enabled.gdb glycan database was selected. The ‘keep top N candidates’ feature was set to 50, and Data Type was set as HCD with child scan dissociation set as EThcD. The ‘maximum Oglycan allowed’ was set to five unless otherwise noted. In silico digestion parameters were set to generate decoy proteins using reversed sequences, and the initiator methionine feature was set to ‘variable’. The maximum modification isoforms allowed was 1,024, and the minimum and maximum peptide length values were set to 5 and 60, respectively. Number of missed cleavages was set to 3. Precursor and product mass tolerances were 10 and 20 ppm, respectively, and the minimum score allowed was three. Modifications were set as carbamidomethyl on C as fixed, oxidation on M, and N-terminal acetylation as a variable. All identified glycopeptide spectra were manually verified for accurate identification.

### Sample Preparation for Native MS

Each unprocessed Fabs, containing 10 μg of the protein, was buffer-exchanged into 150 mM aqueous ultrapure ammonium acetate (pH 7.2) by ultrafiltration (Millipore Amicon) with a 10 kDa cutoff filter. The protein concentration was adjusted to 2–3 μM prior to native mass spectrometry analysis.

### Native MS Analysis

Samples were analyzed using an Orbitrap Ascend Tribrid Mass Spectrometer (Thermo Fisher Scientific) operating in intact protein mode. Samples were ionized using borosilicate nanoelectrospray emitters fabricated in-house from borosilicate capillaries (0.78 mm inner diameter, 1.00 mm outer diameter) (Sutter Instruments). The spray voltage was set to 1.5 kV, source fragmentation was 60 V, source temperature was 300°C, and ions were mass analyzed in the Orbitrap with a resolution set to 30k (at m/z 200). Each Fab was measured for at least 1 min continuous acquisition.

### Native MS Data Analysis

The accurate masses of the Fabs were extracted by averaging and deconvoluting the electrospray ionization spectrum to a zero-charge spectrum using UniDec ^64^. The standard settings were except for an expected peak of 0.85 Th, a point smooth width set to 10, and artifact suppression (Beta) set to 0.

### Deglycosylation of mAbs and SDS-Page Quantification

Mab deglycosylation was performed with Endoglycosidase H (Endo H, NEB #P0702) and Endoglycosidase S (Endo S, NEB #P0741) according to the manufacturer protocol. Briefly, 20 microgram of monoclonal antibody were combined with 10XBuffer and 1 microliter of deglycosylase in total volume of 20 microliter. Reaction was performed at 37oC overnight. Half of the reaction was analyzed on SDS-PAGE for completion of the deglycosylation reaction.

### Antiserum adsorption

Mice were immunized with purified lectin domain of FimH from *E. coli* strain K12. Activity of the obtained antiserum was determined by serial dilutions in ELISA using 96-well plates coated with purified FimH-FocH fimbriae. Dilution factor that is the closest to the saturation level of chromogenic signal (within 1:5,000-30,000 range depending on the day of experiment) was used for the adsorption experiments. Bacteria expressing fimbriae with no FimH and with FimH-FocH or FimH-FocHF1L variants were grown overnight, washed with PBS, concentrated to OD650=9.0, and mixed 1:1 with the diluted antisera in the Eppendorf tube. The adsorption was done by the tube 16 rpm rotation at room T for 1h. Upon the completion, the mixture was centrifuged 10,000 rpm per 3 min, the supernatant was filtered through .45 uM membrane and used in the ELISA.

### Enzyme-linked immunosorbent assay

ELISA for the mAb475 deglycosylation and anti-FimH serum adsorption experiments was performed as described previously ^27^. Briefly, type 1 pili containing the tips identical to those used for the cryo-EM reconstruction (FimH from *E. coli* strain K12) were expressed by and sheared off the recombinant bacteria. Purified pili were immobilized on the surface of microtiter wells. The monoclonal antibodies or polyclonal serum were added for 45 min incubation. The level of antibody binding was determined by anti-mouse IgG secondary antibodies coupled with HRP. The average of means obtained in three independent experiments are plotted on the graphs with combined standard deviation shown as error bars, and P-values calculated in Student’s T-test for pairwise comparison to evaluate the significance independently (command ttesti in STATA v14).

### Sequence Alignments of mAB475-like Antibodies

Outgroup antibody sequences were identified using the Protein BLAST tool from NCBI. The light chain sequence of mAb475 was used as the query. N- and C-terminal amino acids from the mAb475-like antibodies were trimmed. MEGA7 was used to align sequences and build the phylogenetic tree ^65^. After the initial alignment, the tree was trimmed to only include the branches containing at least one mAb475-like antibody.

### Antibody Sequencing (Fab21)

Sequencing of V-Region of mAb Light and Heavy Chains. Total mRNA was isolated from hybridoma cells using an AllPrep® DNA/RNA Mini kit (Qiagen), followed by cDNA synthesis with the iScript™ RT cDNA Synthesis Kit (BioRad) according to manufacturer’s procedure. The V-regions were PCR amplified using 2-3 μL from the 25-μL RT reaction volume. For Light Chain, following primers were used: forward: VL4: CCAGTTCCGAGCTCCAGATGACCCAGTCTCCA; VL5: CCAGATGTGAGCTCGTGATGACCCAGACTCCA; VL7: CCAGTTCCGAGCTCGTGATGACACAGTCTCCA; and reverse: VLr:GCGCCGTCTAGAATTAACACTCATTCCTGTTGAA. For Heavy Chain, following primers were used: forward: VH1: AGGTCCAGCTGCTCGAGTCTGG; VH3: AGGTCCAGCTTCTCGAGTCTGG; VH5: AGGTCCAACTGCTCGAGTCAGG; and reverse: VH3_1: AGGCTTACTAGTACAATCCCTGGGCACAAT; VH3_2a: GTTCTGACTAGTGGGCACTCTGGGCTC. Gel-purified PCR products were subjected to sequencing by Genewiz Inc.

### Molecular Dynamic Simulations

Molecular dynamics (MD) simulations were performed using the complexes determined through cryo EM (FimH-Fab926 and FimH-Fab475). For simplicity and to reduce the computational cost, FimH was truncated after residue 160 (keeping only the lectin domain), while the heavy and the light chain of Fab926 were truncated after residues 115 and 112, respectively (keeping only the respective variable domains). Briefly, the simulations were performed with the program NAMD and the CHARMM22 force field ^66,67^. The protein was solvated using a cubic water box with side length of 118 Å and 125 Å for the complex with mAb926 and mAb475, respectively, and periodic boundary conditions. Chloride and sodium ions were added to neutralize the system and approximate a salt concentration of 150 mM. During the simulations, the temperature was kept constant at 300 K or 330 K by using the Langevin thermostat with a damping coefficient of 1 ps^-1^, while the pressure was held constant at 1 atm by applying a pressure piston ^68,69^. First, a 50-ns long simulation was performed at 300 K in order to determine stable inter-domain side chain contacts between LD and the Fab926 domains and the domains with each other. This was followed by a 50-ns long simulation at 330 K to accelerate sampling of events. To determine whether mannose clashes with the 29-31 loop of the Fab926 heavy chain we superimposed each frame of the trajectories onto a previously published model of lectin domain in the inactive conformation bound to mannose ^26,43^. The superimposition was done by aligning the Cα atoms of residues 1 to 6 and 44 to 48 of the lectin domain followed by visual inspection of the resulting model.

### Rigid-Body Docking

The online tool pyDockWEB ^70^ was used to perform docking of Fab926 to a conformation of LD with bound mannose where the clamp loop is closed, corresponding to the active conformation (PDB code 1UWF) and to a conformation of lectin domain where the clamp loop is open but mannose is still bound, corresponding to a snapshot sampled after 330 ns of a 330-K simulation performed in ^43^. PyDockWEB performs protein-protein rigid-body docking using electrostatics and desolvation energies. As pyDockWEB can only perform protein-protein docking, docking was performed in the absence of mannose.

## Notes

### Summary of Updates

There were errors in the author list. Evgeni Sokurenko's name was not capitalized, so I fixed that. And I changed the corresponding author to me (Justin Kollman) - the three last authors are co-corresponding, but it seems I can only select one author for this, and I will handle the manuscript and updates. Otherwise, the rest of the manuscript is untouched.

