## Supplemental Information for "Antibodies disrupt bacterial adhesion by ligand mimicry and allosteric interference"

**Movies**

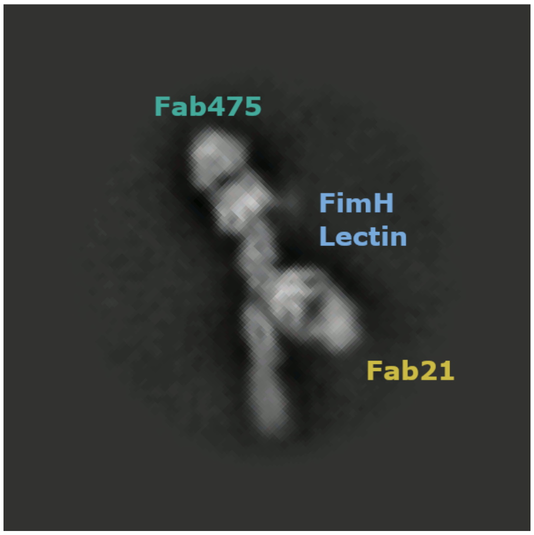


**Movie 1: Series of 2D class averages of Fab21 and Fab475 bound to FimH.** Class averages show the pilin domain and FimG in multiple positions. See Ext. Data Fig. 3e for individual still images.

**Ext. Data Tables**

| mAB/Fab Designation | FimH residues interacting with Fab light chain | FimH residues interacting with Fab heavy chain | Buried Area (Å$^2$) | | | Prior Data Source |
|---|---|---|---|---|---|---|
| | | | Light | Heavy | Total | |
| mAB21 | 23-24, **26**-27, 118, 122, 151-**153**, **155** | 25-**26**, 27-28, **29**-30, 32-33, 35, 37, 117, **155-157**, 158 | 302 | 516 | 756 | [23] |
| mAB475 | **1**, 13, **46**-48, 51, **52, 54**, **133**, **135**, 137, 138, 140, 142 | 51, **52**, 55, 92, 134, **135**,**136**, 137, 138, 139 | 372 | 384 | 712 | [26] |
| mAB824 *FimH inactive conformation* | 55,78,**79-80**,**82**,88-**90**, **91**, 92, 94 | 27, 29-30, 32, **34**, 37, 39-40, 74, 76, 78, **79-82**, 83, 87, 88, 101-102, 104, 108 | 363 | 676 | 983 | [27] |
| mAB824 *FimH active conformation* | 55,78,**79-80**,**82**,88-**90**, **91**, 92, 94 | 33, **35**, 37, 39, 74, 76, 78, **79-80**, 81-83, 87-88, **91**, 104, 108 | 333 | 537 | 839 | [27] |
| mAB926 | 50, 51, **136**, 137 | 1, 2, 10-15, 46-48, 51, **52**, 54, 133, **135**, **137**, **138**, 139, **140**, 142 | 282 | 727 | 983 | [26] |

**Ext. Data Table 1: FimH Epitope Residues.** Epitope residues were determined with the "measure buriedArea" command in ChimeraX. Bolded residues were identified in prior mutational analyses or NMR studies.

**Ext. Data Table 2: Glycan quantification on the Fab475 light chain.** Fab475 was digested, subjected to mass spectrometry analysis, and all identified glycans were quantified. Quantification was done by integrating the area under the curve of the glycopeptide signal. The table presents the resulting glycan quantification.

**Ext. Data Table 3: Quantification of PNGase F-attributed asparagine deamidation.** Each of the Fabs was digested with trypsin or chymotrypsin, and subsequently, N-glycosylation was cleaved using PNGase F, leaving a deamidated asparagine where N-glycosylation had been. This deamidation was quantified using mass spectrometry and presented in the table. PNGase F-attributed deamidation is identified by detecting deamidated asparagines (indicated in the Ndeamidation column) that are part of the N-sequon (indicated in the NSequonDeamidation column), which is the sequence specific for N-glycosylation (NxS/T, where x is not proline).

**Ext. Data Figures**

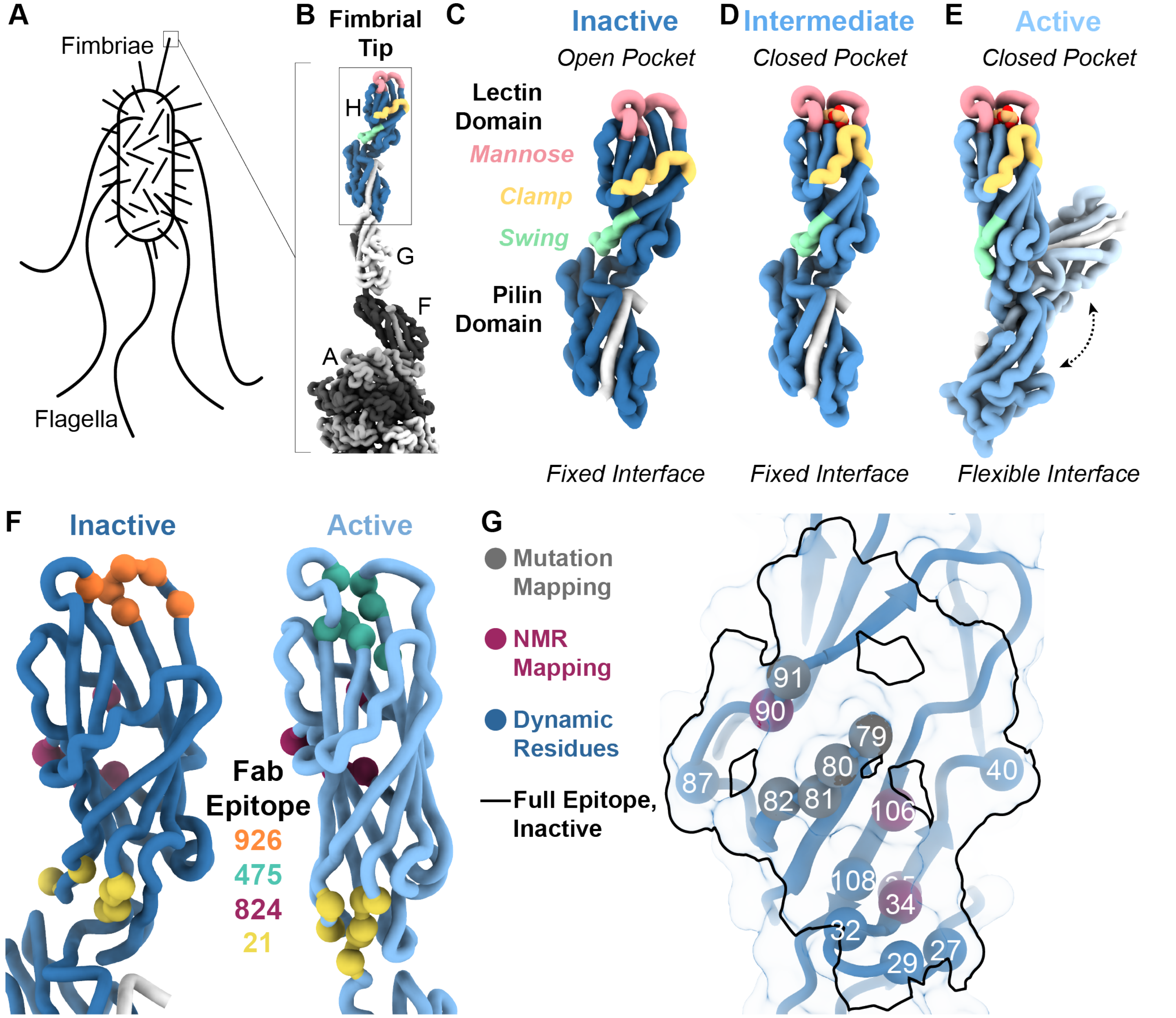


**Ext. Data. Fig. 1: FimH conformations and prior knowledge of antibody epitopes. A**. Cartoon showing flagella and fimbriae of *E. coli*. **B**. Ribbon diagram of the fimbrial tip. FimH in blue, with FimG, F, and A in greys. PDB IDs 6C53, 3JWN. **C-E**. Ribbon diagrams of FimH in the inactive state (**C**, no mannose bound; dark blue; PDB ID 3JWN), an intermediate state with mannose bound (**D**, blue with mannose in orange; PDB ID 4XOE), and the active state (**E**, mannose bound; light blue with mannose in orange; PDB IDs 1KLF, 4XOB). FimH undergoes large conformational changes in the transition from inactive to active. Loops have been highlighted to demonstrate the changes: the swing loop (pale green) and clamp loop (pale gold) move away from each other and the mannose binding loops (pale red) and the clamp loop encircling the ligand. **F.** Cα carbons (spheres) from FimH residues, previously determined by FimH mutagenesis, to interact with bound antibodies mapped onto the inactive (PDB ID 3JWN; dark blue ribbon) and active (PDB ID 1KLF, light blue ribbon) conformations of FimH: orange spheres, Fab926; teal spheres, Fab475; maroon spheres, Fab824; gold spheres, Fab21. **G.** Epitope of Fab824 (black outline) over ribbon representation of FimH + Fab824/926 (transparent surface with blue ribbon). Amino acid Cα of previously identified residues shown as spheres: grey, mutation-identified residues; maroon, NMR-identified residues; blue, dynamic FimH residues.

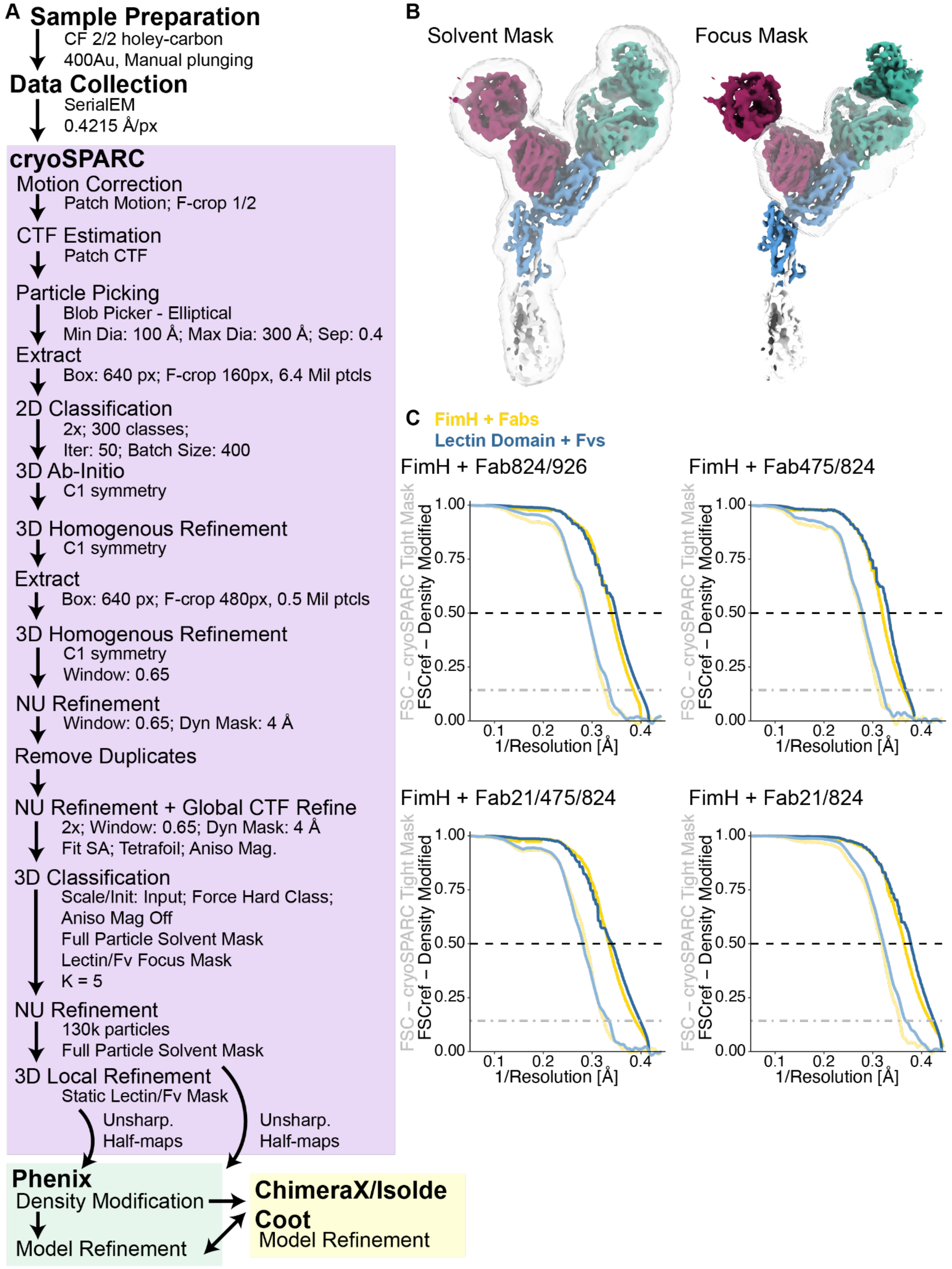


**Ext. Data. Fig. 2**: **CryoEM data processing scheme and data quality. A.** Sample preparation and data collection strategy for FimH-Fab475/824 complex. A similar strategy was adopted for each high-resolution dataset. **B.** Masks used for high-resolution structure determination for FimH-Fab475/824. Similar masks were used for each high-resolution dataset. **C.** Fourier Shell Correlation curves for the final maps generated in cryoSPARC and the density modified maps generated in Phenix. Pale colors represent cryoSPARC gold-standard maps with an FSC cutoff of 0.143 (grey dash-dot line) and dark colors represent density modified maps with an FSC cutoff of 0.5 (black dashed line). Yellow curves represent the full FimH/Fab structures with solvent mask only, while blue curves are reconstructions assembled using the focus mask shown in *B*.

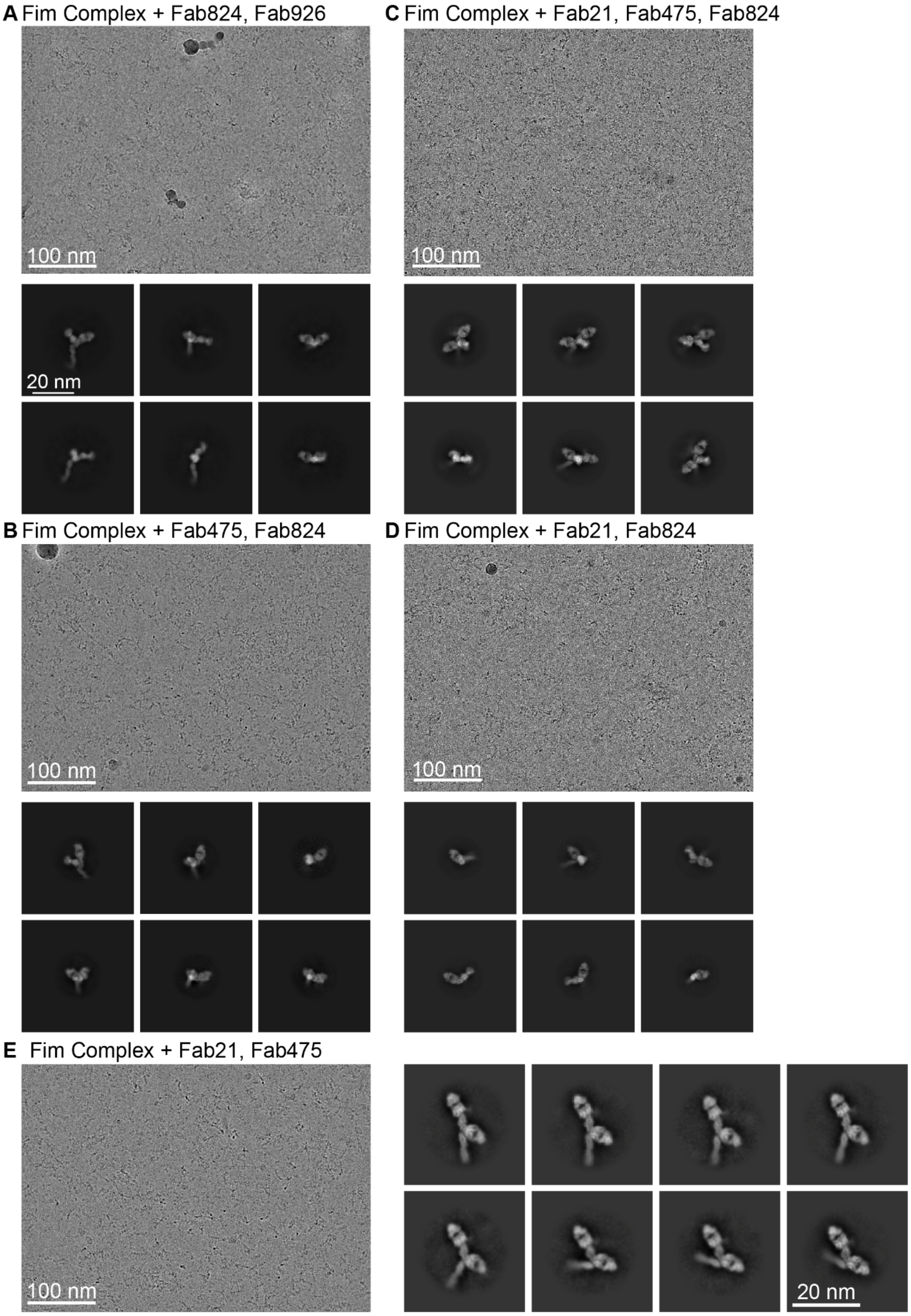


**Ext. Data. Fig. 3: Micrographs and two-dimensional class averages from FimH + Fab datasets. A.** Aligned, dose-weighted, and summed micrograph and two-dimensional class averages from the dataset containing FimH + Fab824/926. **B.** Aligned, dose-weighted, and summed micrograph and two-dimensional class averages from the dataset containing FimH + Fab475/824. **C.** Aligned, dose-weighted, and summed micrograph and two-dimensional class averages from the dataset containing FimH + Fab21/475/824. **D.** Aligned, dose-weighted, and summed micrograph and two-dimensional class averages from the dataset containing FimH + Fab21/824. **E**. Aligned, dose-weighted, and summed micrograph and two-dimensional class averages from the dataset containing FimH + Fab21/475. Two-dimensional class averages have been reoriented and cropped for comparison.

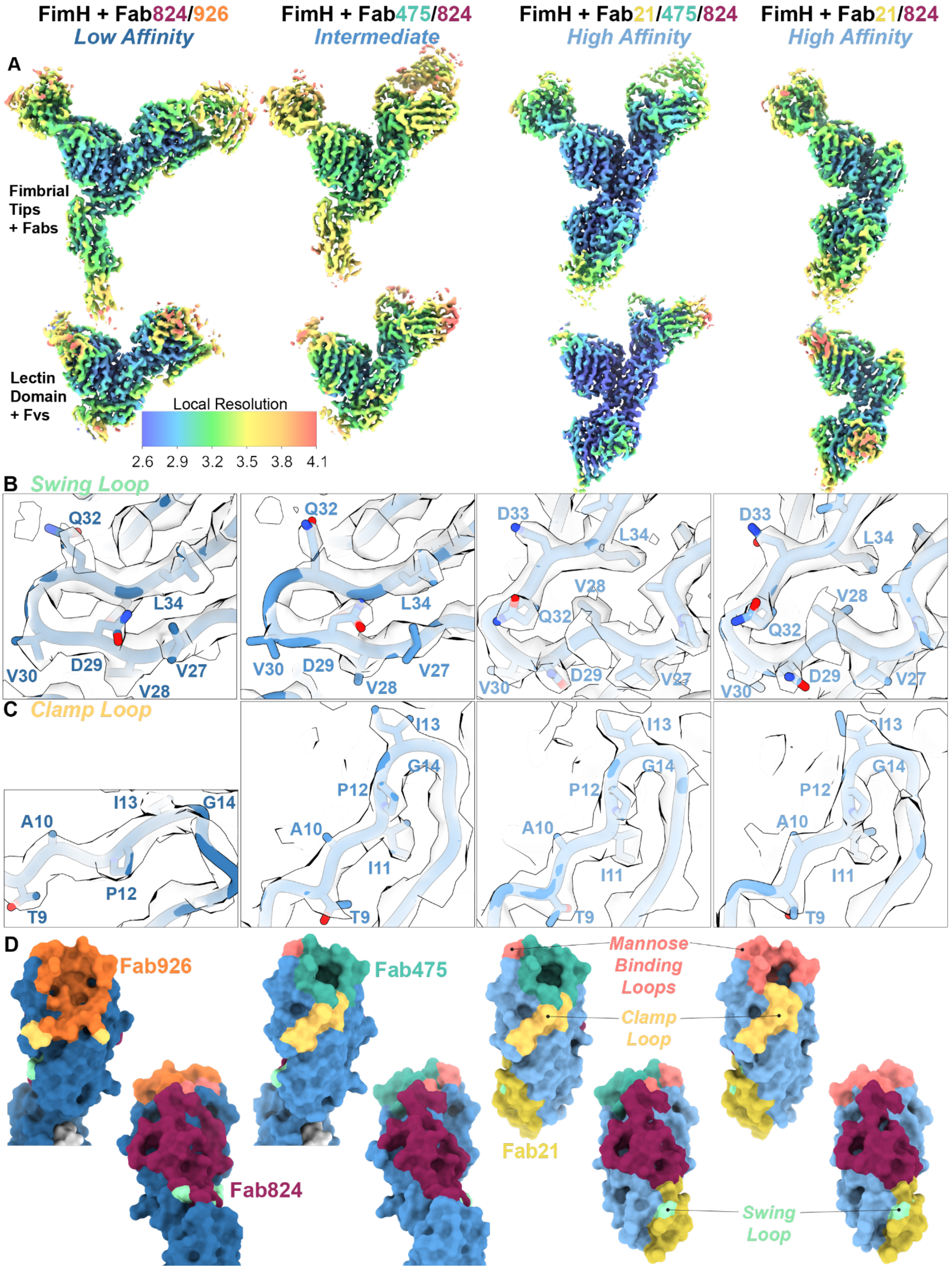
FimH + Fab824/926
Low Affinity
FimH + Fab475/824
Intermediate
FimH + Fab21/475/824
High Affinity
FimH + Fab21/824
High Affinity
A
Fimbrial Tips + Fabs
Lectin Domain + Fvs
Local Resolution
2.6 2.9 3.2 3.5 3.8 4.1
B Swing Loop
Q32 L34 V30 D29 V27 V28
Q32 L34 V30 D29 V27 V28
D33 L34 V28 Q32 V30 D29 V27
D33 L34 V28 Q32 V30 D29 V27
C Clamp Loop
A10 I13 G14 P12 T9
I13 G14 P12 A10 I11 T9
I13 G14 P12 A10 I11 T9
I13 G14 P12 A10 I11 T9
D
Fab926
Fab824
Fab475
Mannose Binding Loops
Clamp Loop
Fab21
Swing Loop

**Ext. Data Fig. 4: CryoEM data quality and FimH epitope mapping. A.** Local resolution estimation for all high resolution structures, calculated in cryoSPARC and mapped onto the density modified volumes; scale shown at bottom for all maps. **B.** Volume and model for the swing loop from each of the high resolution datasets. **C.** Volume and model for the clamp loop from each of the high resolution datasets. **D.** Surface representation of FimH from each of the high resolution structures with epitopes (dark colors) mapped onto the surface. Surfaces are 180° rotated for each model. Fabs: orange, Fab926; teal, Fab475; maroon, Fab824; gold, Fab21. Loops: pink, mannose binding; light yellow, clamp; light green, swing. Quantification of epitope surface area can be found in Ext. Data Table 1.

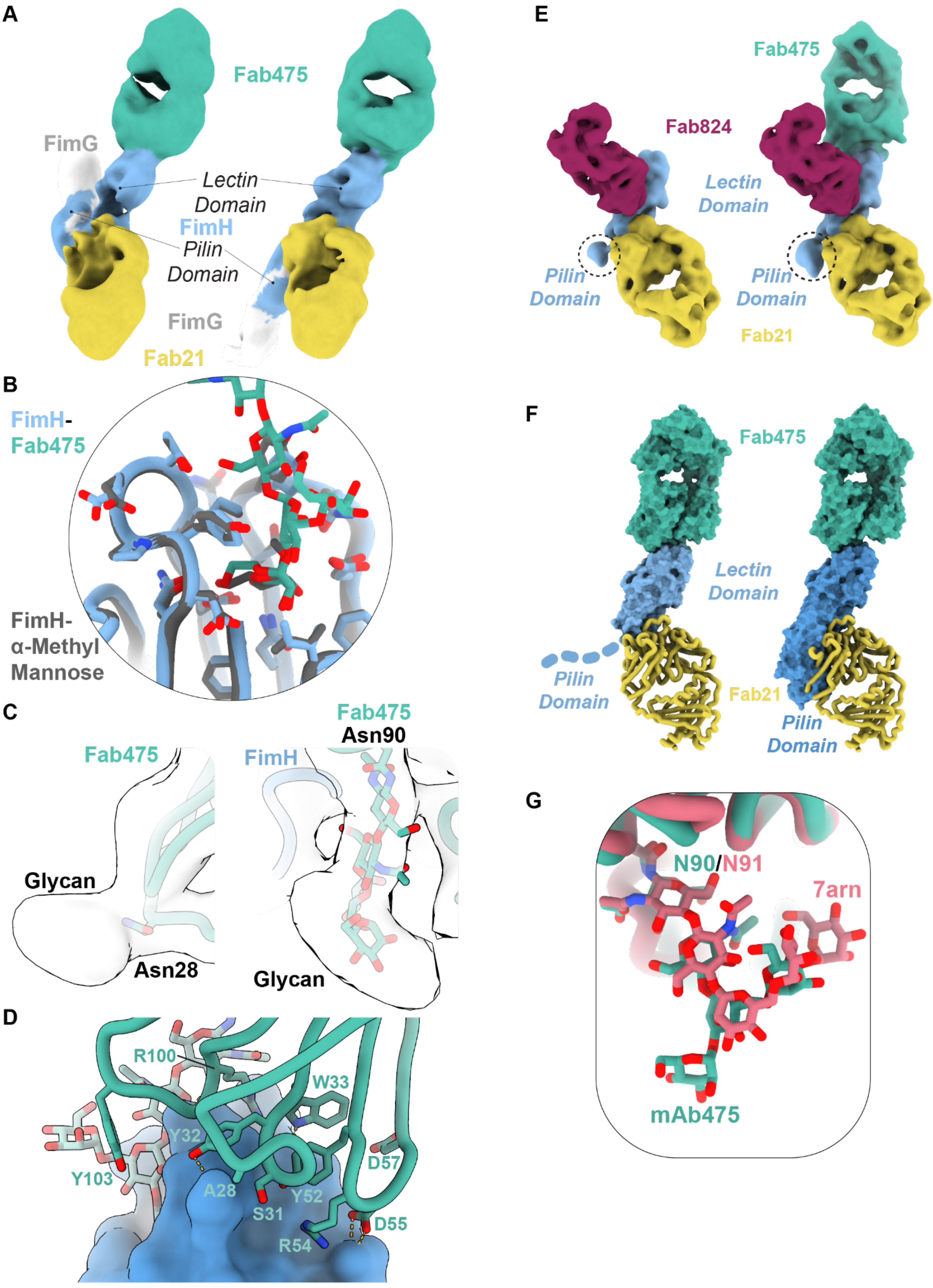
A
Fab475
FimG
Lectin
Domain
FimH
Pilin
Domain
FimG
Fab21
B
FimH-
Fab475
FimH-
α-Methyl
Mannose
C
Fab475
Glycan
Asn28
Fab475
Asn90
FimH
Glycan
D
R100
W33
Y32
Y103
A28
S31
Y52
D57
R54
D55
E
Fab475
Fab824
Lectin
Domain
Pilin
Domain
Pilin
Domain
Fab21
F
Fab475
Lectin
Domain
Pilin
Domain
Fab21
Pilin
Domain
G
N90/N91
7arn
mAb475

**Ext. Data Fig. 5: Supplementary figure panels for FimH in complex with Fab475 and Fab21. A**. Reconstruction of FimH bound to Fab21 (gold) and Fab475 (teal) with the pilin domain (FimH, light blue) and FimG (light grey) in two distinct orientations. **B**. Alignment of FimH lectin domains from the FimH-Fab21/475/824 complex (FimH: light blue sticks and ribbon; Fab475: teal sticks and ribbon) and the FimH-Fab21/824 complex (grey sticks and ribbon). **C**. Low-pass filtered volume of FimH in complex with Fab475 showing volume for two glycosylation sites, Asn28 and Asn90. **D**. Contact surface between Fab475 (teal) and FimH (blue). Fab475 residues within 5 Å of FimH shown. **E**. Volumes of FimH + Fab21/824 (left) or Fab21/475/824 (right) reconstructed with only a large circular mask and gaussian filtered. Portion of pilin domain present in volumes indicated with dashed circle. F. Surface representation of Fab475 (teal) and FimH (blue) demonstrating how Fab21 (gold ribbon) does not clash with the pilin domain in the active conformation (light blue, left) but does clash in the inactive or intermediate conformations (blue, right). **G**. Overlay of glycan from Fab475 (teal) with glycan from a Fab derived from a surface immunoglobulin (PDB ID: 7arn; rose). Alignment on the Fab light chains, with the panel centered on the glycans.

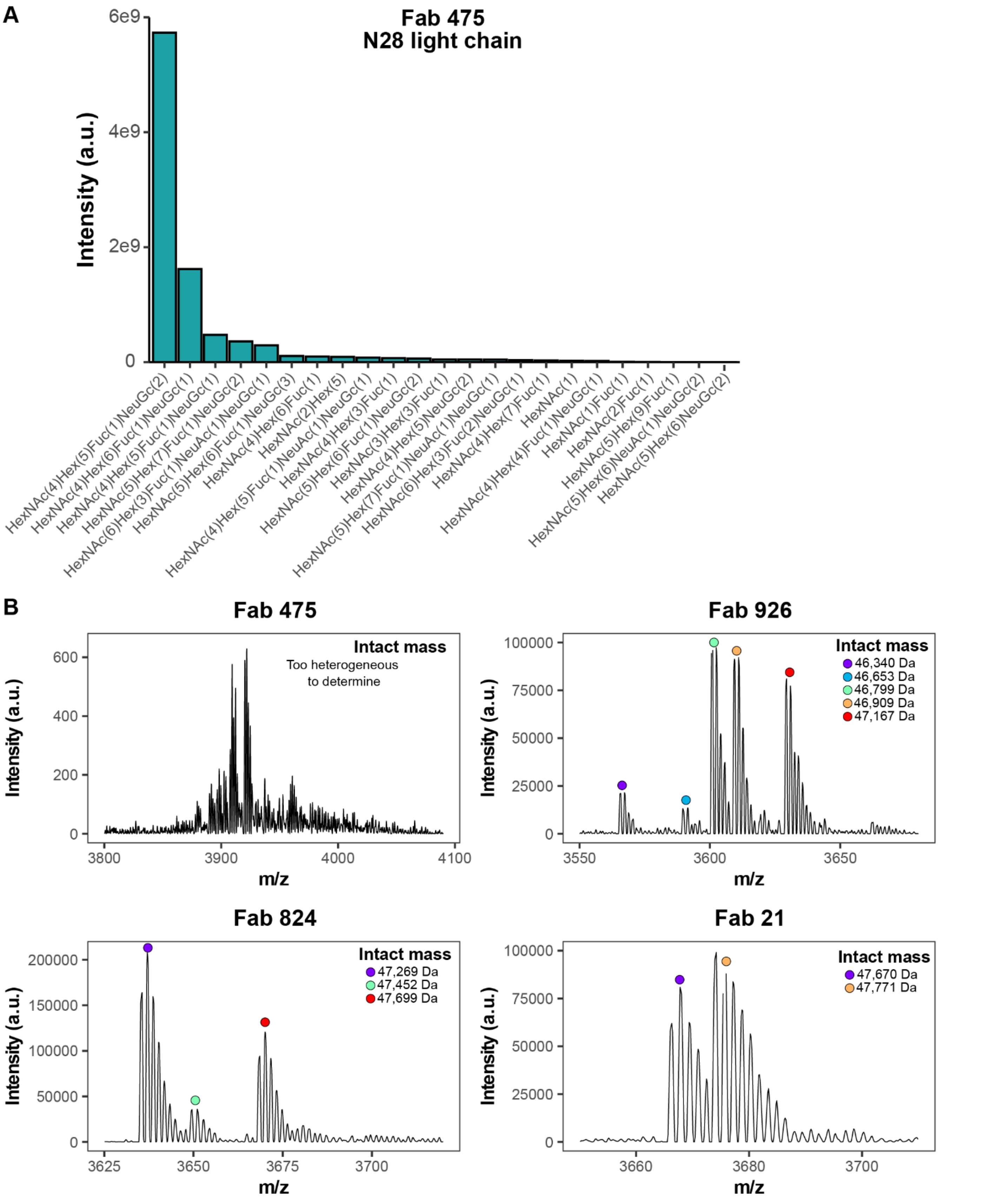


**Ext. Data Fig. 6: Quantification of N-glycosylation 28 on the Fab 475 light chain and intact mass spectrometry analysis of the Fabs.** Fab molecules were subjected to intact mass spectrometry analysis, and the extracted ion chromatogram of one charge state envelope is shown. Fab 475 showed highly heterogeneous spectra preventing intact mass calculations, which is typical for glycoproteins. Fab molecules 926, 824, and 21 were deconvoluted using UniDec [2015MartyetalAnalChem], and their calculated intact mass is presented.

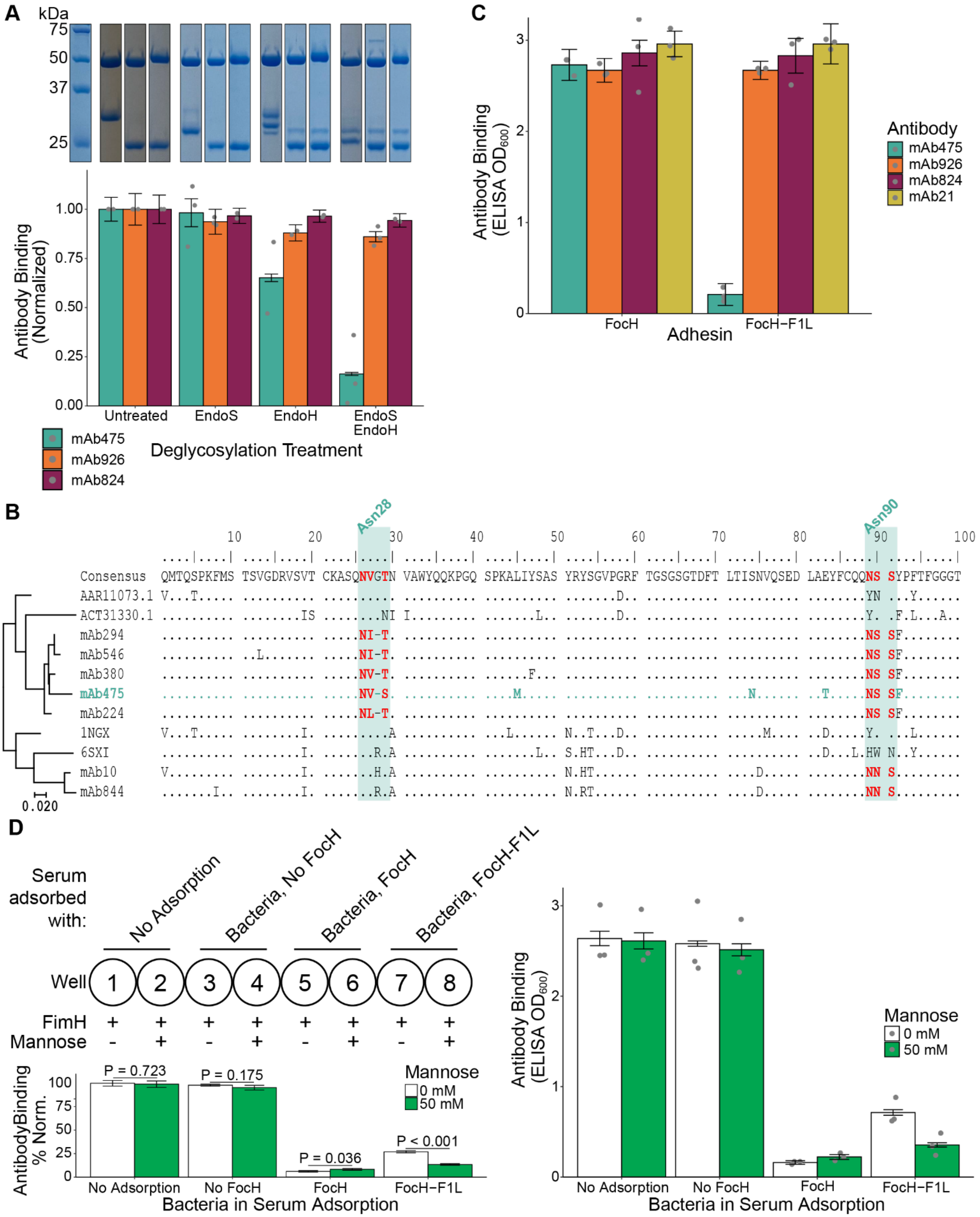

A
kDa
75
50
37
25
Antibody Binding (Normalized)
1.00
0.75
0.50
0.25
0.00
Untreated
EndoS
EndoH
EndoS EndoH
Deglycosylation Treatment
mAb475
mAb926
mAb824
C
Antibody Binding (ELISA $OD_{600}$)
3
2
1
0
FocH
FocH−F1L
Adhesin
Antibody
mAb475
mAb926
mAb824
mAb21
B
Asn28
Asn90
10
20
30
40
50
60
70
80
90
100
Consensus QMTQSPKFMS TSVGDRVSVT CKASQNVGTN VAWYQQKPGQ SPKALIYSAS YRYSGVPGRF TGSGSGTDFT LTISNVQSED LAEYFCQQNS SYPFTFGGGT
AAR11073.1
ACT31330.1
mAb294
mAb546
mAb380
mAb475
mAb224
1NGX
6SXI
mAb10
mAb844
0.020
D
Serum adsorbed with:
No Adsorption
Bacteria, No FocH
Bacteria, FocH
Bacteria, FocH-F1L
Well
1
2
3
4
5
6
7
8
FimH
Mannose
AntibodyBinding % Norm.
100
75
50
25
0
P = 0.723
P = 0.175
P = 0.036
P < 0.001
Mannose
0 mM
50 mM
No Adsorption
No FocH
FocH
FocH−F1L
Bacteria in Serum Adsorption
Antibody Binding (ELISA $OD_{600}$)
3
2
1
0
Mannose
0 mM
50 mM
No Adsorption
No FocH
FocH
FocH−F1L
Bacteria in Serum Adsorption

**Ext. Data Fig. 7: Asn90 glycan on mAb475 is critical to its function and is representative of a pool of antibodies found within the polyclonal antibody response. A**. *Top:* SDS-Page gel evaluation of Fab fragments digested with endoglycosidase S, endoglycosidase H, or both endoglycosidases. Full, uncut acrylamide gels can be found in Supp. Fig. 1. *Bottom*: Normalized ELISA results of antibody after glycosylation treatment listed on the x-axis. mAb475, mAb926: N = 3 biological replicates; mAb824: N = 2 biological replicates. Error bars = combined S.D.. **B**. Sequence alignment of light chains from mAb475, mAb475-like, and outgroup antibodies. mAb475 shown in teal; teal boxes indicate Asn28 and Asn90 glycosylation site locations. Text in red indicates glycosylation motif present (Nxs/T, x ≠ P). Scale bar represents number of substitutions per site. **C**. ELISA results for each antibody in this study against the FimH variant FocH and the mutant FocH-F1L. N = 3 biological replicates. Error bars = S.D. **D**. *Left*: Diagram of serum adsorption and FimH ELISA setup for identification of mAb475-like antibodies in the serum pool; normalized results and significances shown in lower left plot. Error bar = combined S.D.. *Right*: ELISA results in the absence (white) or presence (green) of mannose with serum-adsorbed samples. N = 3 biological replicates. Error bars = S.D.

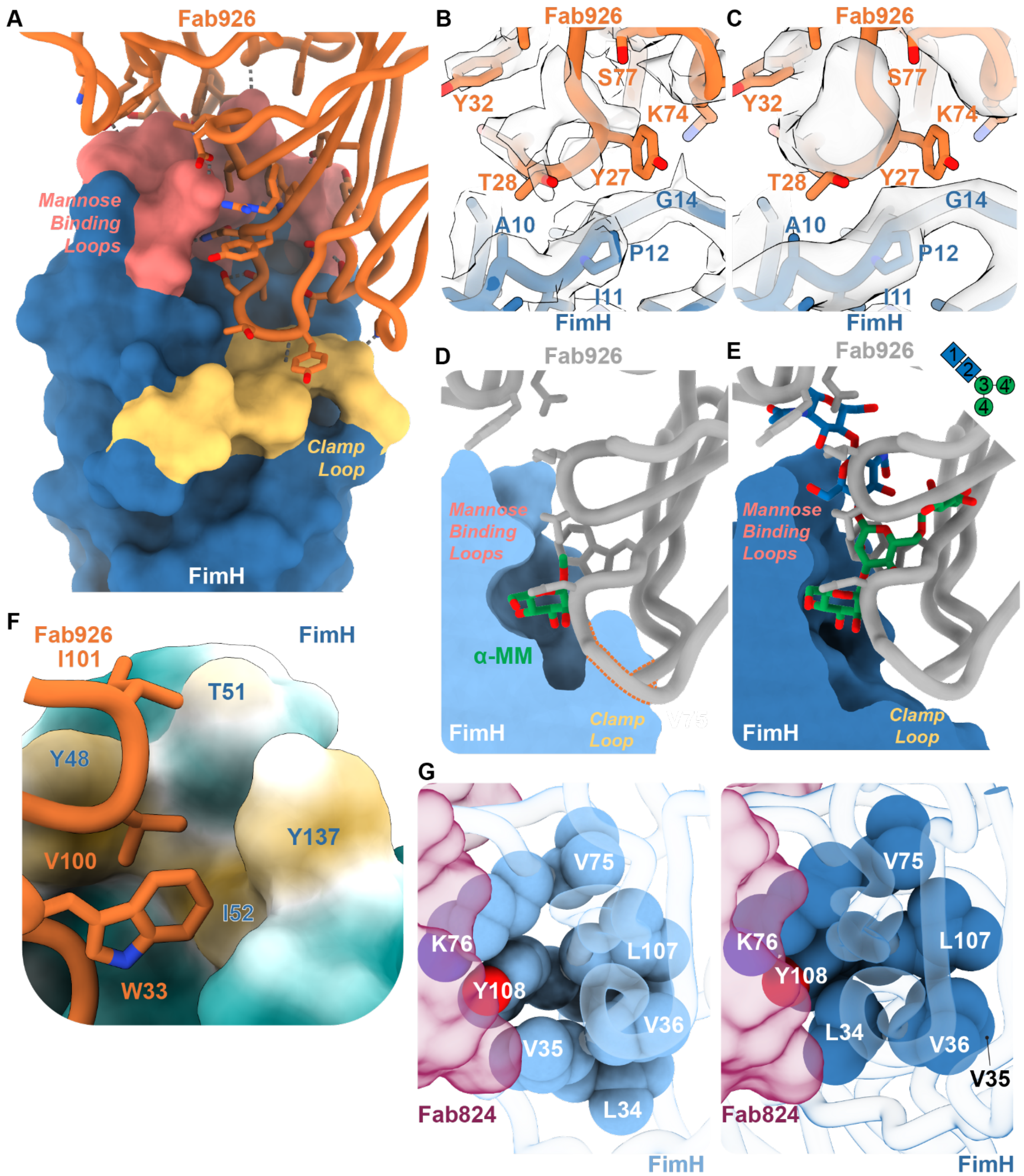


**Ext. Data Fig. 8. Supplementary figure panels for FimH in complex with Fab926 and Fab824. A**. Ribbon and stick diagram of Fab926 showing residues buried in interaction with FimH (dark blue surface). Hydrogen bonds indicated with dashed grey lines. **B-C**. Fab926 (orange) interface with FimH clamp loop (dark blue). *B* shows low-pass filtered volume from *C*. **D**. Alignment of FimH (light blue, surface) in complex with α-methyl mannose (green sticks) with Fab926 (gray sticks and ribbon). Structures were aligned on Cα carbons 1, 46-54, and 136-140 of FimH. Orange dotted lines indicate Fab926 loops that clash with the FimH clamp loop. **E**. Alignment of FimH (dark blue, surface) in complex with Fab926 (gray sticks and ribbon) with oligomannose-3 (green/blue; PDB ID 2vco). Structures were aligned on Cα carbons 1, 46-54, and 136-140 of FimH. **F**. FimH surface showing hydrophobic patch (hydrophobicity shown as yellow to teal transition; yellow: more hydrophobic; teal: more hydrophilic). Fab926 residues shown in orange. **G**. Residues in beta strands near the toggle switch residues L34/V35 (blue spheres) in FimH (transparent blue ribbon) pack tightly against each other and Fab824 (transparent maroon surface).

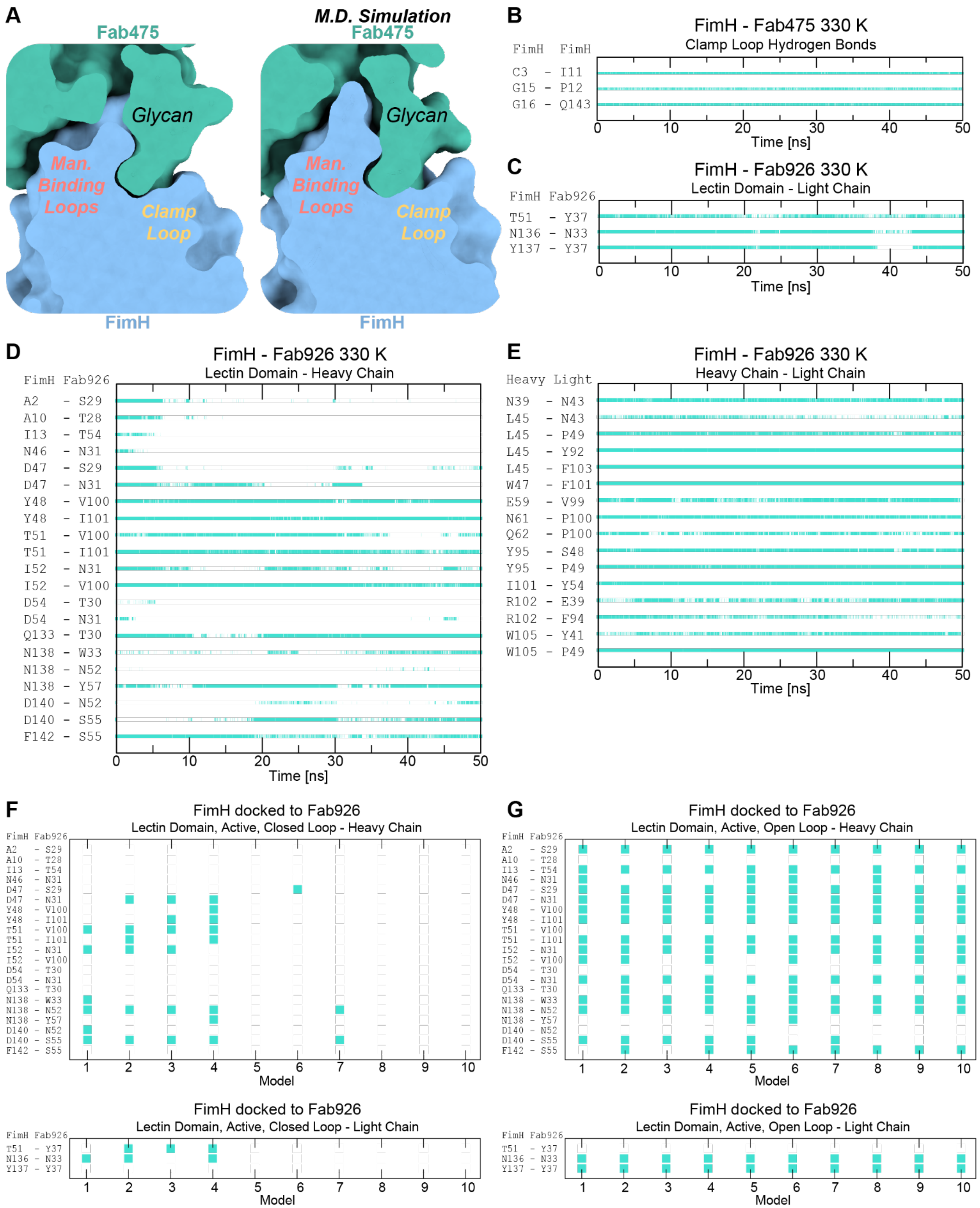


**Ext. Data Fig. 9. Molecular dynamics simulations suggest flexibility at the FimH-Fab926 binding interface. A**. Slice through the cryoEM structure (left) and the model from the molecular dynamics simulation at 50 ns (right); FimH (light blue surface) bound to Fab475 (teal surface). **B**. Time series of hydrogen bond contacts between FimH clamp loop and the rest of FimH. The frames in which a contact is formed in the simulation are indicated by the vertical lines in cyan. **C-D**. Time series of side chain contacts formed between FimH and the variable domains of Fab926. The frames in which a contact is formed in the simulation is indicated by the vertical lines in cyan. **C**. FimH-Fab926, light chain contacts. **D**. FimH-Fab926, heavy chain contacts. **E**. Time series of side chain contacts formed between the two chains of Fab926. The frames in which a contact is formed in the simulation is indicated by the vertical lines in cyan. **F-G**. Models and side chain contacts established from rigid body docking of FimH with Fab926 in pyDockWEB. Cyan squares represent side chain contacts formed. **F**. FimH lectin domain in the active conformation with the clamp loop closed docked to Fab926. **G**. FimH lectin domain in the active conformation with the clamp loop open [from 2016Interlandi&ThomasProteins] docked to Fab926.

**Supplementary Data**

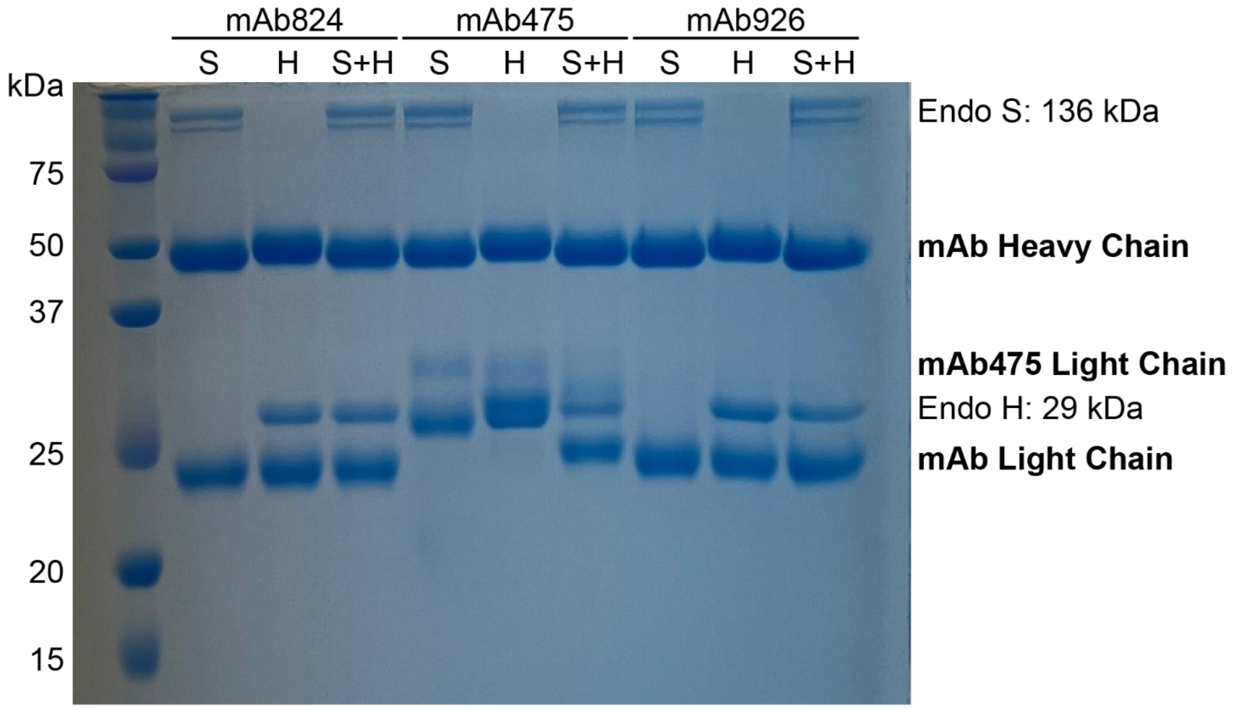


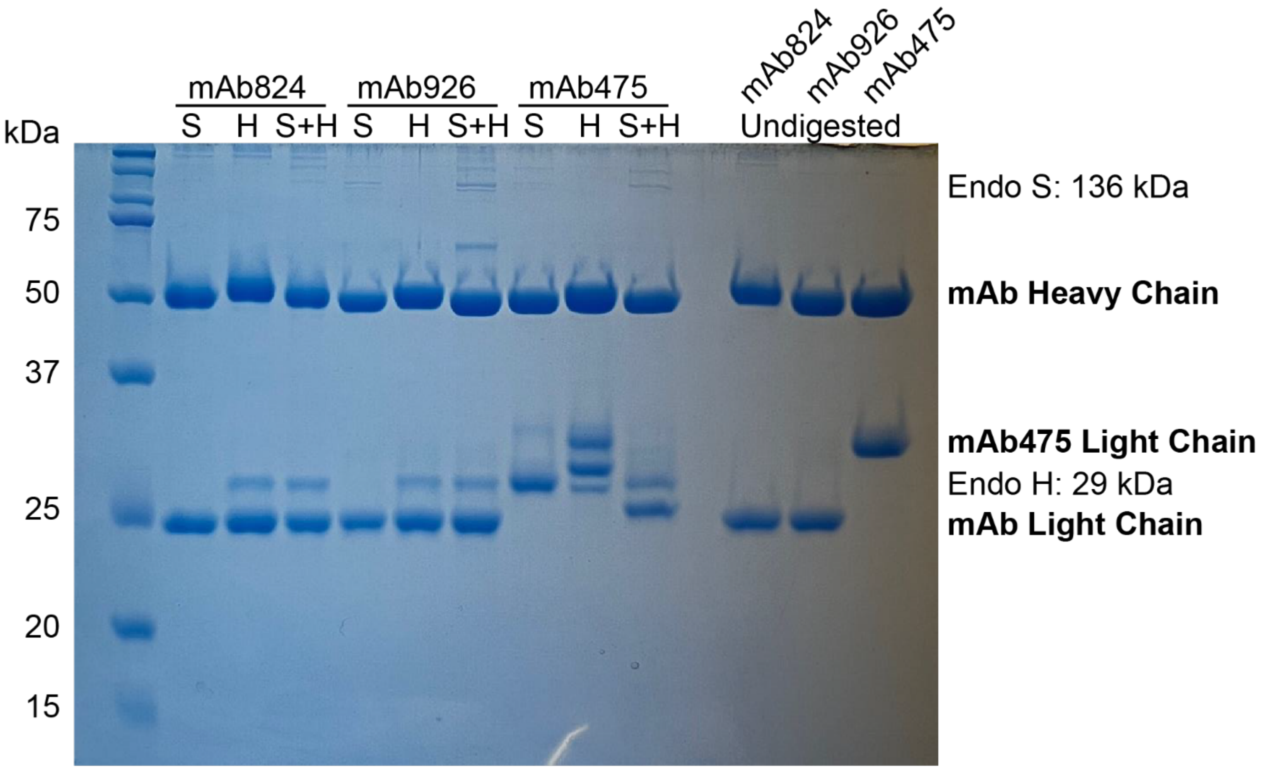


**Supplementary Figure 1. SDS-PAGE gels of endoglycosidase-treated mAbs.** Sections from lower gel shown in Ext. Data Fig. 7.
